# Regulation of mRNA polyadenylation governs mammalian body plan formation in gastruloids

**DOI:** 10.1101/2025.07.22.666114

**Authors:** David Taborsky, Fabiola Valdivia-Francia, Neguin Ranjbar, Laura Llop-Grau, Clara Duré, Umesh Ghoshdastider, Peter F. Renz, Ramona Weber, Merve Yigit, Aleksei Mironov, Katie Hyams, Stefano Vianello, Mihaela Zavolan, Matthias P. Lutolf, Ataman Sendoel

## Abstract

The establishment of the body plan during gastrulation represents a hallmark of animal life. It emerges from the interplay of gene-regulatory programs and positional cues, yet how these signals are integrated post-transcriptionally remains largely unexplored. Here, we combine the scalability of mouse gastruloids with a single-cell CRISPR screening platform to functionally dissect germ layer specification at single-cell transcriptomic resolution. Focusing on post-transcriptional regulation, we systematically map drivers of mesodermal and endodermal fate and identify the deadenylase *Cnot8*. Loss of *Cnot8* leads to widespread poly(A) tail elongation and transcript stabilization, shifting mesoderm differentiation toward ectopic notochord fate, thereby profoundly impacting axial patterning. Collectively, our findings identify mRNA deadenylation as a fundamental mechanism linking cellular identity with morphogenetic signaling during mammalian body plan formation.

## Introduction

The formation of the embryo *in vivo* is an exquisitely complex process, integrating intrinsic and positional cues, coupled with dynamic cellular and tissue reorganization to establish the multidimensional body plan. Central to this process is gastrulation (*1, 2*), the event in which the multipotent and uncommitted epiblast gives rise to the three germ layers, ectoderm, mesoderm and endoderm. Gastrulation has been extensively studied using advanced imaging (*3*) and sequencing tools (*4, 5*), providing insights into the transcriptional regulation of embryonic development, body axis formation and germ layer differentiation across species. Yet, how intrinsic and extrinsic signals are integrated at the post-transcriptional level remains largely unknown.

While these studies have greatly advanced our understanding of early embryogenesis, they have often remained observational, with functional perturbations largely limited to individual genes and requiring labor-intensive knockout embryos or cell-type-specific *in vitro* differentiation protocols (*6, 7*). Furthermore, although transcriptional regulation during early development has been extensively studied, post-transcriptional regulatory processes remain poorly defined, partly due to technical constraints and limited access to mammalian embryos. This gap underscores the need for scalable strategies to explore post-transcriptional regulation in a genome-wide manner to systematically define gene-regulatory networks during early development. By combining multiplexed perturbations with single-cell transcriptomic profiling, single-cell CRISPR screens offer a powerful strategy for uncovering novel regulators of embryonic processes (*8, 9*). Despite their promise, the application of these techniques to the embryonic system has seen limited use due to the technical challenges of delivering perturbation libraries to large numbers of accessible cells, highlighting the need for models that combine high cellular throughput with faithful recapitulation of embryonic cell diversity and morphogenesis.

Gastruloids can recapitulate key elements of mammalian embryogenesis *in vitro* (*10–13*). Upon aggregation of mouse embryonic stem cells and exposure to a 24-hour (h) Wnt agonist pulse at 48 h of development (*10, 12, 14, 15*), gastruloids undergo essential steps of early embryonic development, including the emergence of the three germ layers (*10, 11, 13–15*). Germ layer differentiation is accompanied by structural remodeling and gene expression changes (*10–13, 16*), including processes such as epithelial-to-mesenchymal transition and coordinated cell migration (*16*).

Gastruloids capture crucial aspects of early body plan organization, including symmetry breaking and anterior-posterior axis formation, evidenced by spatially restricted expression of patterning factors such as *Hox* genes (*11, 12*). When cultured under specific conditions - including exposure to an extracellular matrix - gastruloids can undergo morphogenetic processes that give rise to complex structures, including somites (*11, 17*), neural tubes (*17*), and cardiac precursors (*18, 19*). Consequently, gastruloids have enabled extensive profiling of developmental processes, spanning epigenetic (*20*), transcriptomic (*11–14, 21*), metabolomic (*22–25*) and proteomic (*26*) landscapes.

Here, we couple gastruloids with a single-cell CRISPR screening strategy (*8*) to establish a platform for the systematic identification of novel regulators of germ layer differentiation. Focusing on mRNA turnover and translational regulators, we uncover novel factors of germ layer specification in the mesodermal and endodermal lineages. We demonstrate that C*not8*, a deadenylase within the CCR4-NOT complex, orchestrates mammalian body-plan patterning via widespread control of mRNA decay. Our findings reveal a key developmental role of adenylation regulators, beyond the maternal-to-zygotic transition, demonstrating their involvement in governing body plan specification.

## Results

### Establishing a single-cell CRISPR platform in gastruloids

To systematically uncover novel post-transcriptional regulators of germ layer differentiation, we combined the CRISPR droplet sequencing (CROP-seq) strategy (*8*) with gastruloids (*10, 15*). Starting from 406 genes associated with the gene ontology (GO) term “regulation of translation” (GO:0006417), we curated a list of 201 candidate genes based on their expression in gastruloids (Figure S1A). In addition, we included 4 depletion controls and 8 known regulators of germ layer differentiation (herein referred to as germ layer controls). To target these genes, we designed a library containing 669 single guide RNAs (sgRNAs), with 3 sgRNAs per candidate gene, alongside sgRNAs for depletion controls, germ layer controls and 30 nontargeting control sgRNAs. We used a piggyBac-tetCas9 construct to introduce doxycycline-inducible Cas9 into a *Sox1/Brachyury* reporter (SBR) mouse embryonic stem cell (mESC) line proven to be well suited for gastruloid culture (Figure S1B) (*13, 27*). The resulting tetCas9-SBR cells showed normal growth and elongation (Figure S1C). We validated doxycycline-induced Cas9 expression by western blot and confirmed editing efficiency using sgRNAs targeting *Rpl11*, resulting in the expected depletion of infected cells (Figure S1D, E). Following lentiviral production, the sgRNA library was used to infect tetCas9 mESCs on day −10 prior to the initiation of organoid culture. Cas9 expression was induced on day −5 for 72 h using doxycycline and gastruloids were aggregated on day 0 (Figure 1A). Gastruloids were harvested 120 h post-aggregation for single-cell RNA sequencing (scRNA-seq) coupled with sgRNA capture (Figure 1A). We detected a total of 134,727 cells after doublet removal, filtering and sgRNA annotation. To prevent disruption of gastruloid development due to potential depletion of essential genes, we kept the proportion of sgRNA-positive cells low (8.3–8.5%) (Figure 2B). Flow cytometry and time-lapse imaging revealed clearly separated clusters of GFP^+^ cells and a progressive overall depletion of the library by day 5 of gastruloid development (120 h post-aggregation), suggesting an effective depletion of essential translational regulators in the library (Figure 2A-D). In contrast, control guides were evenly distributed across cell types (Figure 1D-F, S2A, B) and showed stable propagation throughout gastruloid development until day 5 (Figure 2C, S2C).

**Figure 1:**
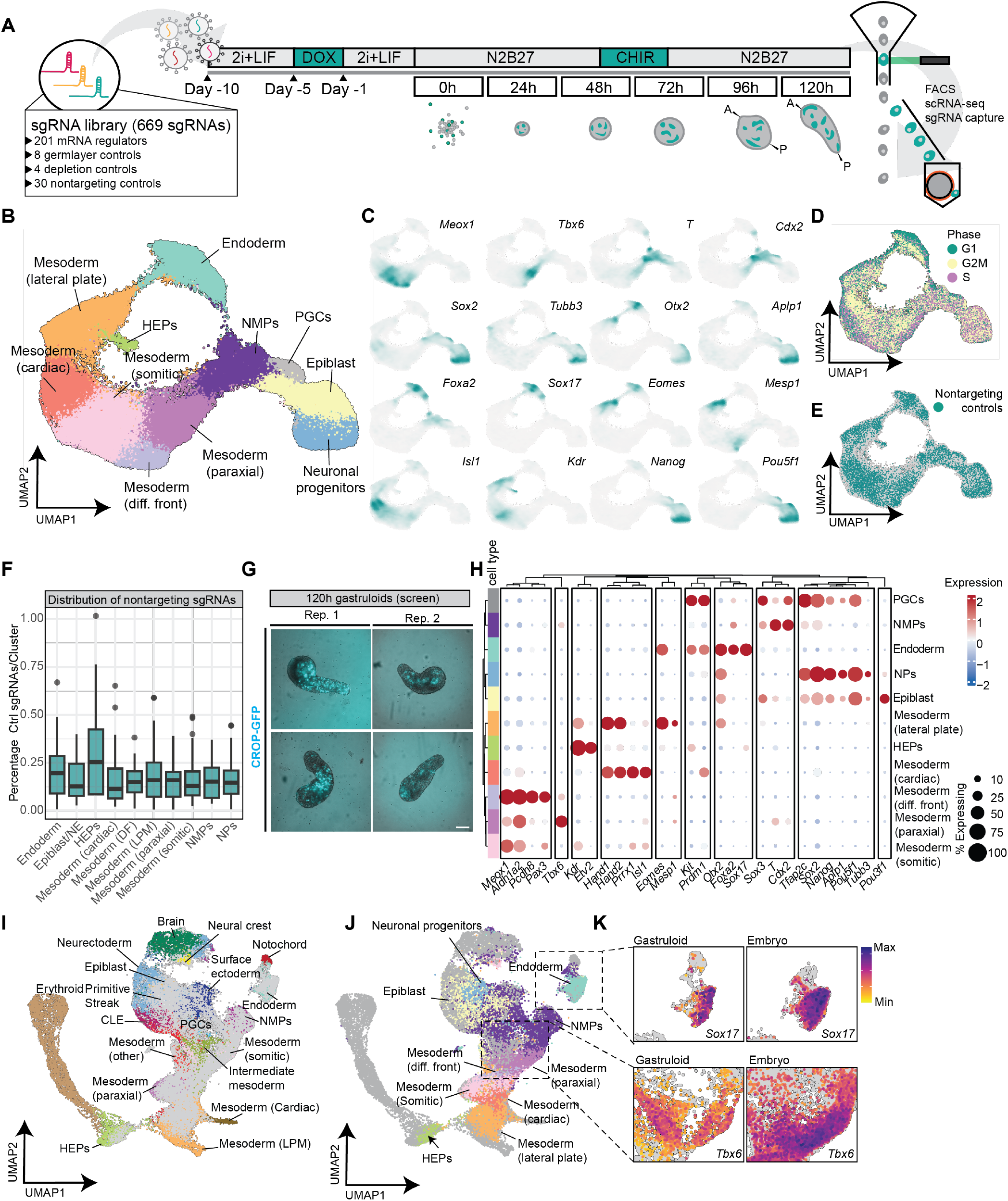
Establishing a single-cell CRISPR platform in gastruloids. **(A)** Schematic of the screen workflow. **(B)** UMAP embedding of 134,727 single cells from two replicate 120h gastruloid CRISPR screens, identifying 11 distinct cell types after quality filtering and sgRNA assignment. **(C)** Marker gene expression profiles across major cell types identified in 120 h gastruloids. Marker sets include mesodermal populations (*Meox1, Tbx6, Mesp1, Isl1*), neuromesodermal progenitors (*T, Sox2, Cdx2*), neuronal progenitors (*Tubb3, Sox2, Aplp1, Otx2*), endoderm (*Sox17, Foxa2, Eomes*), pluripotent epiblast (*Pou5f1, Nanog*) and early hematopoietic progenitors (*Kdr*). **(D-E)** UMAP projections of 120 h gastruloids showing cell cycle states **(D)** and the uniform distribution of the 30 nontargeting control sgRNAs **(E)** across all cell types. **(F)** Box plot depicting the distribution of the 30 nontargeting control sgRNAs across all cell types. **(G)** Representative images of 120h gastruloids in the screen. Scale bar, 200 µM. **(H)** Dot plot showing expression of cell-type-specific markers in the respective cell types. **(I-J)** UMAP projection of 134,727 cells from 120 h gastruloids mapped onto a reference atlas of mouse embryos spanning the developmental stages E7.5-E8.5. Gastruloid cells are shown in grey, with embryonic cell types highlighted **(I)**. Embryonic cell types in grey, with gastruloid clusters highlighted, are shown in **(J). (K)** Feature plots showing the expression of endoderm marker *Sox17* and paraxial mesoderm marker *Tbx6* across gastruloid and mouse embryos.

**Figure 2:**
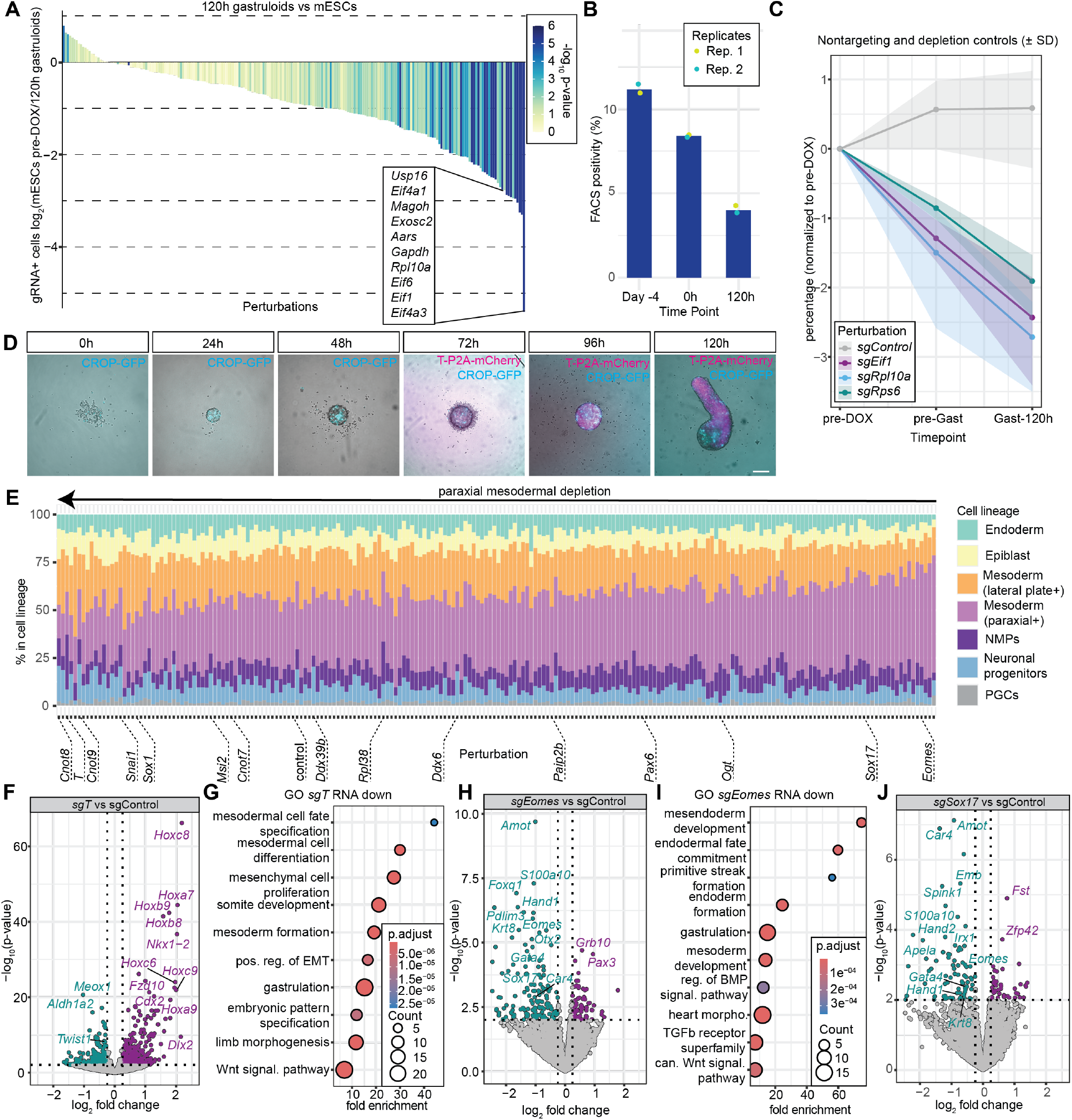
A single-cell CRISPR screen for identifying post-transcriptional regulators of germ layer differentiation in gastruloids. **(A)** Waterfall plot displaying log_2_ fold changes in 120h gastruloid cell numbers relative to initial sgRNA representation in infected mouse embryonic stem cells (mESC) prior to doxycycline induction. A total of 67 target genes are significantly depleted (p < 0.01). **(B)** Percentage of sgRNA-GFP positive cells across the different time points. **(C)** Stable propagation of the nontargeting control library. Median abundance of 30 nontargeting control sgRNAs compared to the depletion of *Eif1, Rps6 and Rpl10a* (3 sgRNAs per gene) across 3 time points, pre-DOX induction (preDOX), pre-gastruloid aggregation (preGast) and after 120h of gastruloid development (WTA). **(D)** Time-lapse imaging of developing gastruloids from 0h (mESCs) to 120h, showing sgRNA-GFP expression and T-P2A-mCherry emerging from 72 h post-aggregation onward. **(E)** Cell type distribution plot of the 213 perturbations and nontargeting controls, sorted by perturbations most depleted in paraxial mesoderm. **(F)** Volcano plot displaying the top differentially expressed genes in 120 h gastruloids upon perturbation of *T* (*sgT*). **(G)** GO term enrichment of genes downregulated (p-value < 0.01) in all *sgT* cells. **(H)** Volcano plot displaying the top differentially expressed genes in 120 h gastruloids upon perturbation of *Eomes* (*sgEomes*). **(I)** GO term enrichment of genes downregulated (p-value < 0.01) in all *sgEomes* cells. **(J)** Volcano plot displaying the top differentially expressed genes in 120 h gastruloids upon perturbation of *Sox17* (*sgSox17*).

We identified cells belonging to all 3 germ layers in our dataset, as well as undifferentiated epiblast cells. Gastruloids showed an overall mesodermal bias with 81,423 cells (60% of total cells) belonging to various mesodermal lineages, including somitic (high in *Meox1*), paraxial (*Tbx6*) and lateral plate/cardiac mesoderm (*Hand1, Hand2, Isl1*) (Figure 1B, C, H). Additionally, we identified an endodermal cluster with high expression of *Sox17* and *Foxa2* and an ectodermal cluster high in *Tubb3* and *Aplp1*, as well as neuromesodermal progenitors (NMPs) high in *Sox2, T/Brachyury* and *Cdx2* (Figure 1B, C, H). Undifferentiated epiblast cells retained expression of the pluripotency markers *Nanog* and *Pou5f1* (Figure 1B, C, H), in line with previous reports showing the persistence of undifferentiated epiblast throughout gastruloid development (*13*). We also observed a minor cluster high in the hematoendothelial progenitor markers *Kdr* and *Etv2* (Figure 1B, C, H) and primordial germ cells (PGCs) high in *Tfap2c* and *Kit* (Figure 1H). Integration and co-embedding of gastruloid cells with embryonic cell data from embryonic day (E) 7.5, E7.75, E8.0 and E8.5 embryos (*4*) showed co-clustering of gastruloid cells with their assigned embryonic cell types, supporting the presence of cell types consistent with *in vivo* embryonic development (Figure 1I-K).

To validate our CRISPR screening strategy and Cas9 efficacy, we first examined known depletion controls. As expected, we found a strong depletion of *Eif1, Rps6* or *Rpl10a* perturbed cells across gastruloid development (Figure 2A, C). We then assessed cell-type-specific depletion phenotypes by comparing the distribution of perturbed cells across different cell types relative to the nontargeting control library (Figure 2E and 3B-F). sgRNAs targeting the germ layer controls *Eomes* and *Sox17* showed endodermal depletion (Figure 2E and 3B), while *T/Brachyury* and *Snai1* sgRNAs were depleted in mesodermal lineages (Figure 2E), validating the high specificity of our screening strategy.

Unlike conventional CRISPR screens, the single-cell CRISPR approach also enables the evaluation of the transcriptomic changes of perturbed cells, providing insights into underlying mechanisms driving the observed phenotypes (*8, 28, 29*). As validation, we first analyzed the transcriptomes of our germ layer controls (Figure 2F-J, S2D-H). *T/Brachyury* is a key transcription factor during gastrulation (*30–32*), particularly in mesodermal lineage commitment. As expected, its perturbation led to downregulation of mesodermal markers including *Meox1* (Figure 2F). This was further supported by GO term enrichment among downregulated genes, which highlighted processes related to gastrulation and mesodermal development (Figure 2G). In line with these observations, depletion of *Eomes*, a transcription factor essential for endoderm induction (*30, 33, 34*), showed a prominent downregulation of genes associated with endodermal development (*Gata4, Sox17, Krt8*) (Figure 2H-I). *Sox17* perturbation similarly resulted in reduced expression of endodermal markers (*Eomes, Gata4, Krt8*) and markers of endocardial differentiation and heart development, including *Hand1* and *Irx1*, consistent with *Sox17’*s essential role in these processes (Figure 2J, S2D) (*35–37*). Furthermore, *Snai1* perturbed cells showed reduced expression of genes associated with mesodermal differentiation (*Mest*) and epithelial-to-mesenchymal transition (*38, 39*) (*Twist1*), while upregulating markers of neuronal lineages (*Sox2*) and undifferentiated cells (*Utf1*) (Figure S2E-F).

### Loss of *Cnot8* results in mesodermal differentiation defects

To investigate germ layer-specific enrichment and depletion of sgRNAs, we used MAGeCK (*40*) to calculate the significance of sgRNA alterations within individual germ layers compared to all other cell clusters. By focusing on germ layer-specific factors that were not globally depleted across gastruloids (Figure 3D), we identified a set of strong candidates with putative roles in post-transcriptional control of germ layer differentiation (Figure 3H-M, S3C-M). These included the RNA-binding protein *Msi2*, which showed marked depletion in the endoderm (Figure 3D). Transcriptomic profiling reflected this phenotype, with downregulation of endodermal cell markers (*Sox17, Eomes*) as well as genes associated with mesoderm and heart morphogenesis (*Tbx6, Mesp1*) (Figure S3D-E). Concomitantly, genes associated with pluripotency (*Dppa5a, Zfp42*) were upregulated, suggesting profound differentiation defects in *Msi2* perturbed cells. Another candidate was the Poly(A) binding protein interacting protein *Paip2b*, whose corresponding sgRNAs showed specific depletion in the NMP population (Figure 3D-E). Transcriptomic analysis of *Paip2b*-perturbed cells supported this phenotype, revealing downregulation of NMP markers such as *Hoxc8* and upregulation of endodermal markers including *Gata4* (Figure S3F-G).

**Figure 3:**
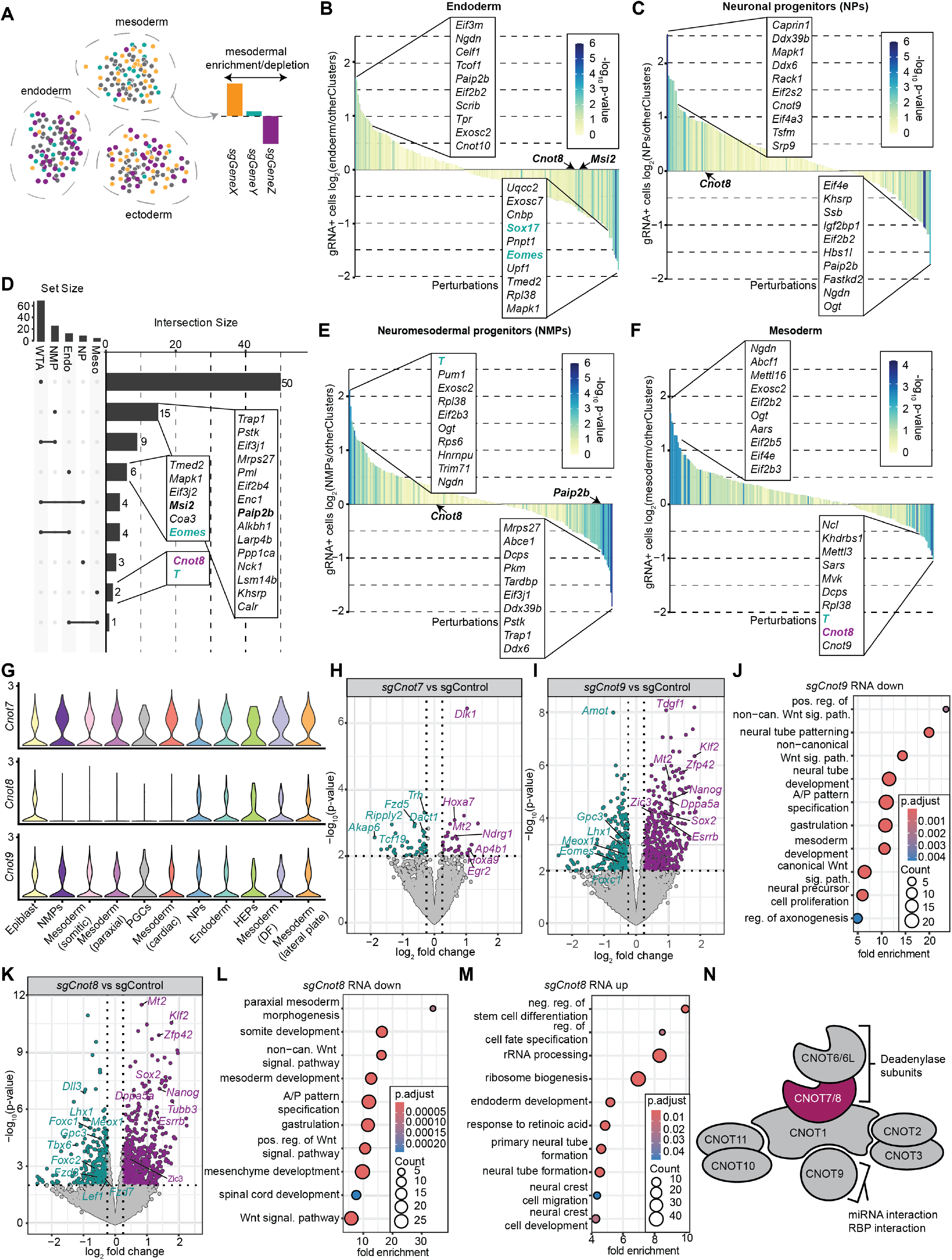
Uncovering novel post-transcriptional regulators of germ layer differentiation. **(A)** Schematic illustrating the strategy for identifying regulators of germ layer differentiation based on cell-type-specific sgRNA enrichment or depletion. **(B-C)** Waterfall plots visualizing changes in sgRNA representation in endoderm **(B)** and neuronal progenitors **(C). (D)** Upset plot displaying depleted targets across all cells (WTA), mesoderm (Meso), endoderm (Endo), epiblast (Epi), and neuronal progenitors (NPs). *Cnot8* and the germ layer control *T* are exclusively depleted in mesoderm (p-value < 0.01, calculated using MaGECK). **(E)** Waterfall plot showing sgRNA enrichment and depletion in neuromesodermal progenitors (NMP). (F) Waterfall plot showing sgRNA enrichment and depletion in mesodermal cells, with *Cnot8* emerging as the most significantly depleted perturbation in mesoderm. **(G)** Violin plot showing the expression of the CCR4-NOT components *Cnot7, Cnot8* and *Cnot9* across gastruloid cell types. **(H-I)** Volcano plot displaying the top differentially expressed genes in 120 h gastruloids upon perturbation of *Cnot7* (*sgCnot7*) and *Cnot9* (*sgCnot9*). **(J)** GO term enrichment of genes downregulated (p-value < 0.01) in all *sgCnot9* cells, showing enrichment for genes associated with non-canonical Wnt signaling and neural differentiation. **(K)** Volcano plot displaying the top differentially expressed genes in 120 h gastruloids upon perturbation of *Cnot8* (*sgCnot8*). **(L)** GO term enrichment of genes downregulated (p-value < 0.01) in all *sgCnot8* cells, showing enrichment for genes associated with paraxial mesoderm development, anterior/posterior (A/P) pattern specification and Wnt signaling. **(M)** GO term enrichment of genes upregulated (p-value < 0.01) in all *sgCnot8* cells, showing enrichment for genes associated with negative regulation of differentiation, maintenance of stem cell populations, neural tube and neural crest differentiation and endoderm formation. **(N)** Schematic of the CCR4-NOT complex with the paralogs CNOT7 and CNOT8 highlighted in magenta.

Intriguingly, CCR4-NOT complex components *Cnot8* and *Cnot9* showed a profound depletion of their corresponding sgRNAs in the mesoderm (Figure 3F), with *Cnot8* being the more significantly depleted candidate, while remaining largely unaltered in overall sgRNA representation and within the endodermal, neuromesodermal and neural compartments (Figure 3B, C, F). This phenotype was more pronounced in paraxial mesodermal lineages compared to the lateral plate mesoderm and its derivatives, suggesting a role of *Cnot8* in this differentiation pathway (Figure 2E and S3A-B).

*Cnot7* is a paralogue of *Cnot8*, with largely similar structure and function but higher binding affinity to the CCR4-NOT complex (Figure 3N, S3N) (*41*). Notably, our single-cell data suggested ubiquitous expression of *Cnot7* across all cell types, while *Cnot8* expression in NMPs and paraxial mesoderm was markedly reduced (Figure 3G). As suggested by the lack of germ-layer-specific depletion of *Cnot7*, perturbation of *Cnot7* had little effect on the transcriptome compared to nontargeting controls (Figure 3H). In contrast, perturbation of another CCR4-NOT component, *Cnot9*, led to strong transcriptomic changes, including downregulation of genes involved in Wnt signaling, gastrulation and differentiation, alongside upregulation of undifferentiated cell markers (Figure 3I-J).

These transcriptomic changes were even more pronounced in *Cnot8* perturbed cells, with marked downregulation of genes associated with mesodermal differentiation (*Meox1, Eomes, Tbx6*), body patterning (*Dll1*) and Wnt signaling (*Lef1*) (Figure 3K). In contrast, markers for neuronal differentiation (*Tubb3*) and pluripotent cell types (*Klf2, Nanog*) showed a robust upregulation (Figure 3K-M). GO term enrichment analysis for genes depleted in *Cnot8* perturbed cells revealed an association with mesenchyme development, somite development, anterior-posterior (A/P) pattern specification, gastrulation and Wnt signaling (Figure 3L). In contrast, genes upregulated in *Cnot8* perturbed cells revealed an enrichment for negative regulators of differentiation, endoderm development and neural tube and crest differentiation (Figure 3M).

We observed a partial overlap between differentially expressed genes upon *Cnot8* and *Cnot9* perturbation, such as the upregulation of pluripotency factors *Zfp42, Klf2* and *Esrrb* (Figure 3I, K, S3O). However, given its more pronounced paraxial mesodermal depletion (Figure S3A) and transcriptomic impact (Figure S3O), we subsequently focused on *Cnot8* and its function in gastruloid differentiation and patterning. Considering the central role of Wnt signaling in NMP differentiation (*42, 43*), promoting mesodermal fate upon activation and neuronal fate upon inhibition (*44, 45*), we surmised that the deregulation of Wnt signaling upon *Cnot8* perturbation may underlie the differentiation phenotypes observed in the screen. Despite their structural similarity (Figure S3N), *Cnot7* and *Cnot8* exhibit distinct functional roles, as evidenced by the lack of mesodermal depletion upon *Cnot7* perturbation, mirroring the viability of *Cnot7* knockout mice compared to the gastrulation-stage lethality observed in *Cnot8* knockouts (*46*).

Consistent with previous reports(*46*), *Cnot8*-perturbed cells showed upregulation in pluripotency markers, including *Klf2, Pou5f1* and *Nanog* (Figure 3K-M). However, in contrast to earlier findings, our data indicated that *Cnot8*-perturbed cells retain the capacity to transition from naïve to formative and primed pluripotent states and can potentially acquire neuromesodermal, neuronal and endodermal fates, while being selectively depleted from the (paraxial) mesodermal lineage. These findings suggest a key role of *Cnot8*-mediated clearance of transcripts beyond the transition from naïve to formative pluripotency, aligning more closely with the *in vivo* phenotypes observed in *Cnot8* knockout embryos (*46*). These embryos generally appear normal until E6.5, forming egg cylinders phenotypically similar to wild-type mice, but then fail during subsequent gastrulation between E7.5-8.5. Since these events take place after the naïve-to-formative transition (*47*), the pronounced developmental defects upon *Cnot8* loss likely reflect disruptions in gastrulation-related signaling and mesoderm commitment.

### *Cnot8* knockout gastruloids show profound patterning defects with induction of notochord cells

To validate the differentiation defects observed in *Cnot8* perturbed cells, we next generated *Cnot8* and *Cnot7* clonal knockout cell lines. We verified the knockout alleles in the clonal cell lines by employing the sequencing-based TIDE assay (*48*) (Figure S4A-H) and evaluated their capacity to form gastruloids (Figure 4A, S4I). Consistent with the results of the single-cell CRISPR screen, while *Cnot7* knockout gastruloids developed normally, *Cnot8* knockout gastruloids showed profound phenotypes (Figure 4A, S4I). Using the *T*/*Brachyury*-P2A-mCherry reporter system in the SBR cell line, we observed that *Cnot8* knockouts failed to generate a T/Brachyury-positive posterior pole and exhibited impaired elongation (Figure 4A). Instead, *T/Brachyury* expression was initially broad at 120 h and later coalesced into multiple poles (Figure 4A, E). While *T/Brachyury* expression decreased over time in both control and *Cnot7* knockout organoids due to NMP depletion, *Cnot8* knockout organoids maintained a robust *T/ Brachyury*-P2A-mCherry^high^ population beyond 144 h of development, confirmed by both time-lapse imaging (Figure 4A) and flow cytometric analysis (Figure 4B-D).

**Figure 4:**
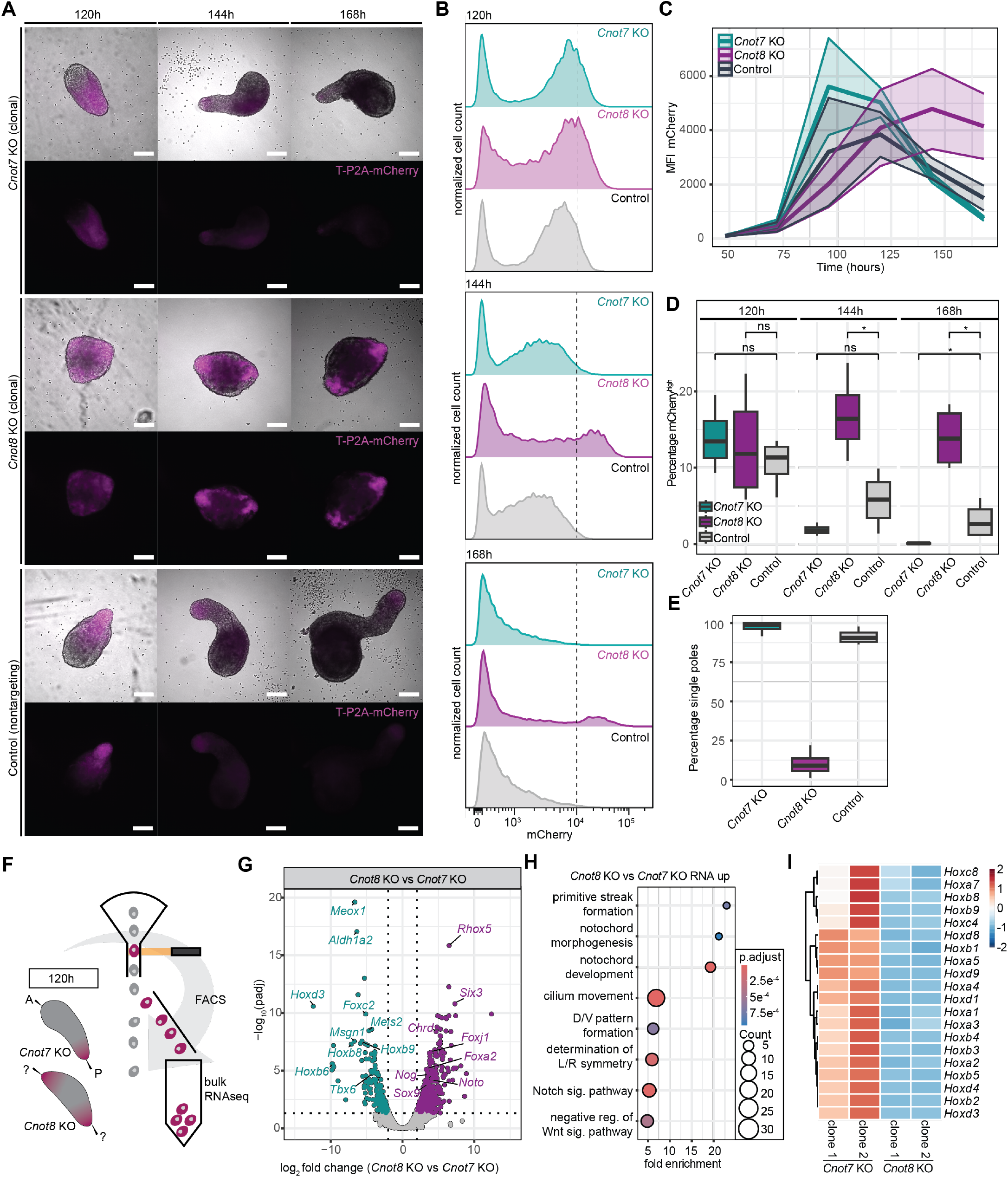
Loss of *Cnot8* deficiency results in gastruloid polarity disruption, higher T expression and appearance of notochord cells. **(A)** Time-lapse imaging of *Cnot8* knockout (KO), *Cnot7* KO and control (nontargeting sgRNA) gastruloids from 120 h (left), 144 h (middle) and 168 h (right), with T-P2A-mCherry expression in magenta. **(B)** Flow cytometry of T-P2A-mCherry in *Cnot8* KO, *Cnot7* KO and control gastruloids from 120 h (top), 144 h (middle) and 168 h (bottom) showing the appearance of an mCherry^high^ population in *Cnot8* KO (dotted line). **(C)** Quantification of the mean fluorescence intensity for *Cnot8* KO, *Cnot7* KO and control gastruloids (n=4 independent gastruloid cultures, 16-8 gastruloids per time point and condition). **(D)** Quantification of the percentage mCherry^high^ cells as indicated in **(B)** (dotted line). Statistics were calculated using a Wilcoxon test. **(E)** Quantification of the number of organoids with single vs multiple poles in 144 h *Cnot8* KO, *Cnot7* KO and control gastruloids (n=4 independent gastruloid cultures, 48 organoids per replicate/condition). **(F)** Schematic of the sorting strategy used to isolate mCherry^high^ cells from 120 h gastruloids for bulk RNA-sequencing. **(G)** Vulcano plot showing differentially expressed genes between *Cnot8* KO and *Cnot7* KO mCherry^high^ cells. **(H)** GO term enrichment analysis of genes upregulated *Cnot8* KO vs *Cnot7* KO mCherry^high^ cells. **(I)** Heatmap showing expression levels of *Hox* genes in *Cnot8* KO vs *Cnot7* KO mCherry^high^ cells from two different cell lines per condition. * p < 0.05

To further investigate this phenotype, we isolated the *T-P2A-mCherry*^*high*^ populations from 120 h *Cnot7* and *Cnot8* knockout gastruloids for transcriptomic profiling (Figure 4F). Differential expression analysis of *Cnot8* knockouts revealed a strong enrichment for GO terms associated with notochord development, including canonical notochord markers like *Foxa2, Noto* and *Chrd*, along with genes associated with cilium movement (Figure 4G-H). These data suggest that the *T/Brachyury*-P2A-mCherry^high^ cells observed in *Cnot8* knockout gastruloids represent notochord cells and not NMPs. Additionally, markers of other mesodermal lineages, especially somitic and paraxial mesoderm markers like *Meox1* and *Tbx6*, were drastically downregulated in *Cnot8* knockouts (Figure 4G), alongside an array of *Hox* genes (Figure 4I), suggesting a loss of anterior-posterior polarity in the gastruloids (*11, 12*). This phenotype was particularly notable as mammalian gastruloids are unable to form notochord cells under standard culture conditions and require special media conditions and signaling cues to differentiate into this specific cell type (*19, 49*).

The notochord is a defining patterning hub in chordate embryos, essential for the formation of the dorsal-ventral body axis and the differentiation of adjacent tissues, including the neural tube, endoderm and paraxial mesoderm (*49*). It is derived from the anterior primitive streak, which gives rise to both axial and paraxial mesoderm (*45*). While the precise mechanisms of notochord formation remain incompletely understood, modulation of Nodal/Activin and more broadly TGFβ signaling (*49, 50*) is required for *in vitro* generation of notochord-like cells. Thus, the formation of notochord cells in *Cnot8* knockout gastruloids points to a substantial signaling landscape shift toward a state permissive for notochord differentiation, ultimately leading to the emergence of a secondary signaling center capable of producing key morphogens such as sonic hedgehog (*Shh*) and bone morphogenetic protein (BMP) inhibitors including *Chrd* and *Nog*.

### CNOT8 regulates transcript poly(A) tail length and transcript abundance

Having confirmed the *Cnot8* perturbation phenotype from the initial screen, we next investigated the molecular mechanisms underlying the observed developmental changes. Given the canonical function of CNOT8 as deadenylase subunit of the CCR4-NOT complex (Figure 3N) (*46, 51*), we examined how its loss impacts the polyadenylation landscape of gastruloids. To accurately quantify changes in poly(A) tail length upon *Cnot8* loss, we performed Nanopore direct RNA sequencing on 72 h gastruloids from *Cnot8* knockout and control conditions (Figure 5A). As expected, *Cnot8* knockout gastruloids showed a marked global increase in median poly(A) tail length (Figure 5B) across all transcripts (Figure 5C-D). Mitochondrial transcripts exhibited no increase in poly(A) tail length, consistent with the known cytoplasmic and nuclear restriction of the CCR4-NOT complex (Figure S6B) (*51*). To identify functionally relevant targets of CNOT8-mediated deadenylation, we assessed candidate transcripts with poly(A) tail changes in *Cnot8* knockouts. Consistent with previous findings (*46*), *Pou5f1, Nanog* and *Dppa5a* showed increased poly(A) tail lengths (Figure 5E, I, S6C-D), supporting a role for CNOT8 in modulating pluripotency. Interestingly, *Zic3* and *Nodal* also showed increased poly(A) tail length in *Cnot8* knockout (Figure 5F-H). *Nodal*, a TGFβ family member, plays an essential role in patterning the early embryo and promoting primitive streak formation (*52, 53*). *Zic3* has been implicated in the formation of axial mesoderm/notochord in other chordates like *Xenopus laevis* (*54*), making both genes promising candidate mediators of the *Cnot8* knockout phenotype.

**Figure 5:**
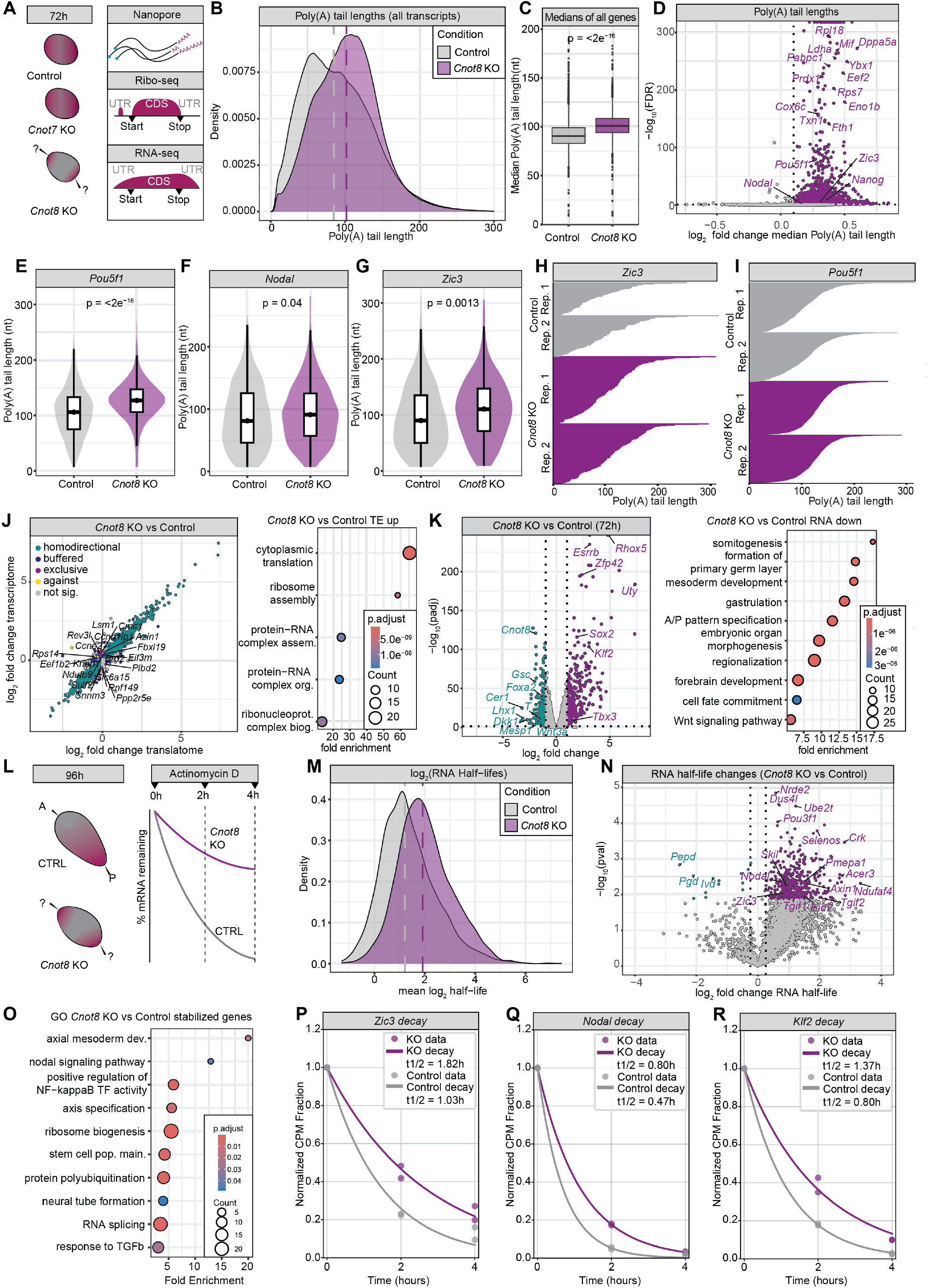
Loss of *Cnot8* results in poly(A) tail elongation and transcript stabilization. **(A)** Experimental outline to dissect *Cnot8* function. Gastruloids from *Cnot8* KO, *Cnot7* KO and control organoids were harvested at 72 h and analyzed by ribosome profiling, bulk RNA-sequencing, and Nanopore direct RNA-sequencing to quantify poly(A) length. **(B)** Density plot showing poly(A) tail lengths across all transcripts in *Cnot8* KO and control 72 h gastruloids. **(C)** Box plot showing the median poly(A) tail lengths in *Cnot8* KO and control gastruloids. **(D)** Volcano plot showing the log_2_ fold change in poly(A) tail length upon *Cnot8*. P values indicate a Wilcoxon test with multiple testing correction using the Benjamini-Hochberg method. (E-G) Box plot showing poly(A) tail lengths of *Pou5f1* (E), *Nodal* (F) and *Zic3* (G). **(H-I)** Plot depicting the poly(A) tail lengths of *Zic3* **(H)** and *Pou5f1* **(I). (J)** Scatter plot showing the log_2_ fold changes in transcript abundance and ribosome occupancy (translatome) of 72h *Cnot8* KO versus control organoids. Genes regulated exclusively at the translational level are shown in magenta. Right panel: GO terms enriched among genes with increased translational efficiency (TE). **(K)** Volcano plot showing differentially expressed genes (transcriptome) in *Cnot8* KO versus control. Right panel: GO term enrichment of transcriptionally downregulated genes in *Cnot8* KO versus control at 72h. **(L)** Schematic of RNA decay experiment. 96 h gastruloids were treated with Actinomycin D and RNA was collected at 0 h, 2 h and 4 h post-treatment. **(M)** Density plot comparing RNA half-lives for *Cnot8* KO and control organoids. **(N)** Volcano plot highlighting genes with altered RNA half-lives in *Cnot8* KO versus control organoids. **(O)** GO terms enrichment of stabilized genes in *Cnot8* KO, suggesting an enrichment for genes associated with axial mesoderm development, Nodal signaling, NF-κB activity and ribosome biogenesis. **(P-R)** Transcript decay plots of stabilized genes in *Cnot8* KO, including *Zic3* **(P)**, *Nodal* **(Q)** and *Klf2* **(R)**.

Deadenylation plays a central role in reducing transcript stability and, in early development, additionally mediates translational repression (*55, 56*). To distinguish between these two outcomes, we first set out to establish a translational landscape of wild-type gastruloids. We performed ribosome profiling from gastruloids across 4 developmental stages, including 48 h, 72 h, 96 h and 120 h post-aggregation. The resulting dataset showed hallmarks of high-quality ribosome profiling datasets, including triplet periodicity (Figure S5A), P-site distribution biased toward coding sequence (CDS) (Figure S5B) and an enrichment of 30-nucleotide footprint sizes (Figure S5C). To assess translational control during development, we examined changes in translational efficiency (TE, defined as ribosome profiling reads divided by RNA sequencing reads) throughout organoid development. Comparing 72 h to 48 h post-aggregation, corresponding to early gastrulation, we observed translational regulation of the translational machinery, including various components of the ribosome and eukaryotic initiation factors (Figure S5D-F). Components of the Wnt signaling pathway also showed altered translational efficiency, consistent with translational buffering of this key developmental pathway in gastrulation (Figure S5E-F). GO term analysis highlighted enrichment of Wnt signaling among genes with reduced translational efficiency (Figure S5F). At later stages, fewer TE changes were detected, which may reflect the limitations of bulk ribosome profiling in resolving translational differences across diverse cell types of late-stage gastruloids (Figure S5G-H).

Having established the translational landscape of gastruloids, we investigated translational changes in *Cnot8* knockouts. We subjected 72 h *Cnot8* knockout, *Cnot7* knockout and control gastruloids to ribosome profiling. *Cnot8* knockout gastruloids displayed widespread transcriptional changes, with 5,451 genes showing differential expression compared to controls (Figure 5J-K, S5I-L). Downregulated genes were enriched for terms related to gastrulation, differentiation and Wnt signaling, while pluripotency-associated genes such as *Sox2, Esrrb*, and *Klf2* were strongly upregulated (Figure 5K). However, when contrasting transcriptomic and translatomic changes, we observed minimal alterations in translational efficiency (Figure 5J), suggesting that *Cnot8* primarily impacts transcript stability rather than translation. In line with the lack of phenotype observed in *Cnot7* knockout organoids, we found minimal transcriptional and translational changes in *Cnot7* knockouts (Figure S5L). Altogether, our integrated 72 h dataset demonstrates that *Cnot8* knockout leads to extensive remodeling of the transcriptome and poly(A) tail landscape, while having a limited impact on the translational efficiency.

### Loss of *Cnot8* increases transcript stability in gastruloids

We next explored the CCR4-NOT complex’s canonical role in transcript destabilization via deadenylation (*51*). To capture a more mature time point of gastruloid development, we performed an RNA stability assay at 96 h post-aggregation. Gastruloids were treated with Actinomycin D to globally inhibit transcription and transcript abundance was quantified at 0, 2 and 4 hours by RNA sequencing (Figure 5L). We first assessed transcriptome-wide differences between *Cnot8* knockout and control gastruloids at 96 h (corresponding to 0 h Actinomycin D time point). We observed widespread transcriptomic changes, including a dramatic loss of Wnt signaling components and regulators, indicative of global repression of Wnt signaling (Figure S6E-F). Additionally, markers of anterior-posterior axis formation such as *Hox* genes and genes essential for mesodermal differentiation, like *Msgn1*, were depleted (Figure S6E). Conversely, markers of neuronal fates such as *Pou3f1* and *Tubb3* were enriched, alongside increased expression of notochordal markers such as *Foxa2, Chrd* and *Tppp3* (Figure S6E).

We then determined the impact of *Cnot8* loss on transcript stability. Consistent with its role in deadenylation, *Cnot8* knockout led to a pronounced increase in transcript half-lives, with median half-lives increasing across hundreds of genes (Figure 5M-N). Interestingly, *Nodal* and *Zic3* wereamong the transcripts with increased stability (Figure 5P-R), consistent with the observed poly(A) tail elongation in *Cnot8*-deficient gastruloids and implicating *Cnot8* in modulating key developmental regulators through mRNA decay.

In addition, we observed transcript stabilization of *Axin1, Eomes* and *Fgf8*, all of which have roles in the formation, patterning and migration of the primitive streak and axial mesoderm (Figure S6G-O) (*57, 58*). Stabilization of *Axin1* (*58, 59*) may underlie at least partly the global downregulation of Wnt signaling, while stabilized *Nodal* and *Fgf8* could indicate activation of TGFβ and FGF signaling. Notably, TGFβ activity may be further influenced by changes in the stability of multiple TGFβ regulators, including repressors such as *Skil, Eid2* and *Tgif2* all of which were stabilized in *Cnot8* knockout gastruloids (Figure 5N, S6J-M). Given that temporally and spatially regulated TGFβ is critical for the differentiation of notochord in embryo models and embryos (*49, 53, 60*), the effect of *Cnot8* knockout on many TGFβ-related transcripts may underlie the ectopic notochord differentiation seen in *Cnot8* knockout gastruloids. In line with this notion, *Zic3* has also been strongly implicated in node formation, migration (*61, 62*) and notochord specification in *Xenopus* (*54*) and mice (*61*).

### Loss of *Cnot8* induces notochord formation and reprograms axial patterning

Finally, we sought to characterize the overall changes in cell type composition in *Cnot8* knockout, *Cnot7* knockout and control organoids using scRNA-seq. We derived *Cnot8* knockout and control gastruloids from two different clonal knockout cell lines harboring different sgRNAs, complemented with gastruloids from a single *Cnot7* knockout clonal cell line. For each condition, we harvested 16 gastruloids, pooled in equal ratios and subjected to scRNA-seq. After removal of doublets, low-quality and dead cells, 16,966 cells were retained for analysis, consisting of 4,873 *Cnot7* knockout, 4,858 *Cnot8* knockout, and 7,235 control cells. Unbiased clustering allowed for the identification of 11 clusters, including mesodermal cells, neuromesodermal progenitors (NMPs), neuronal progenitors, epiblast, endoderm as well as a surprisingly high number of notochord cells (Figure 6A). Marker gene expression was consistent with these cell type assignments, including *Meox1* and *Tbx6* for mesoderm, *Tubb3* and *Otx2* for neuronal progenitors, *T/Brachyury* and *Cdx2* for NMPs, *Sox17* and *Gata6* for endoderm and *Foxa2, Noto* and *Chrd* for notochord (Figure 6A, 6C-D).

**Figure 6:**
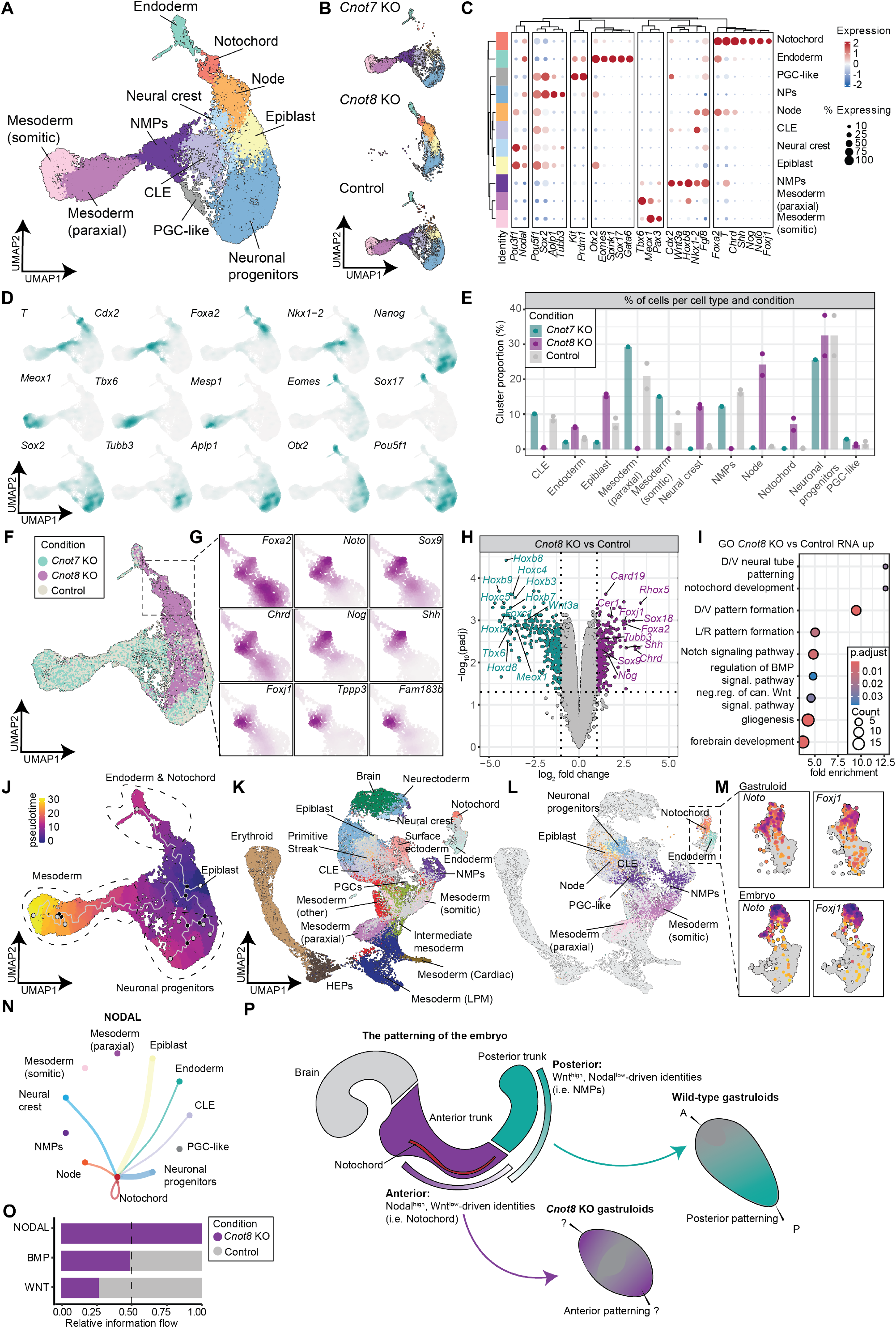
*Cnot8* KO single-cell RNA sequencing reveals altered differentiation trajectories. **(A)** UMAP of 16,966 cells from *Cnot8* KO, *Cnot7* KO and control (nontargeting sgRNA) gastruloids at 120h post-aggregation. **(B)** Individual UMAPs of *Cnot7* KO (top), *Cnot8* KO (middle) and control (bottom) gastruloids. **(C)** Dot plot showing marker gene expression across the identified cell types. **(D)** Nebulosa density plots showing cell-type markers for neuromesodermal progenitors (NMPs) (*T, Cdx2, Nkx1-2*), mesoderm (*Meox1, Tbx6, Mesp1*), endoderm (*Sox17, Eomes, Otx2, Foxa2*), neuronal progenitors (NPs) (*Tubb3, Aplp1, Sox2, Otx2*), epiblast (*Pou5f1, Nanog*), node and caudal lateral epiblast (CLE) (*Nkx1-2*) and notochord (*Foxa2*). **(E)** Bar plot quantifying cell type composition in *Cnot7* KO, *Cnot8* KO and control gastruloids. **(F)** UMAP highlighting *Cnot8* KO enrichment in notochord and anterior primitive streak/node regions. **(G)** Nebulosa density plots showing the expression of canonical notochord markers (*Foxa2, Noto, Sox9, Chrd, Nog, Shh, Foxj1, Tppp3* and *Fam183b*). (H) Volcano plot displaying the top differentially expressed genes in *Cnot8* knockout vs control gastruloids across all cell types. **(I)** GO term enrichment of genes downregulated (p-adj < 0.05, log_2_ fold change < −1) in all *Cnot8* knockout cells. **(J)** Pseudotime trajectory analysis comparing differentiation dynamics in *Cnot7* KO, *Cnot8* KO and control organoids. (K-L) UMAP showing the mapping of gastruloid cells to a reference embryo atlas (E7.5–E8.5). **(K)** UMAP showing embryonic cell clusters with gastruloid cells in grey, **(L)** UMAP showing gastruloid cell clusters with embryonic cells in grey. **(M)** Feature plot of *Noto* and *Foxj1* expression in gastruloids (top) and embryonic (bottom) cells. **(N)** CellChat analysis of NODAL signaling in *Cnot8* KO gastruloids, showing high activity between notochord and other cell types. **(O)** Relative information flow of NODAL, BMP and WNT signaling between *Cnot8* KO and control gastruloids. **(P)** Proposed model for *Cnot8*-mediated body plan formation. Deletion of *Cnot8* prolongs the half-life of transcripts that specify anterior identity (*Nodal, Axin1, Zic3*), shifting developmental fates toward primitive-streak and anterior cell fates, including notochord. This stabilization indirectly suppresses Wnt pathway activity, leading to diminished expression of posterior markers (*Hox, Cdx*) and impaired posterior axis formation.

The emergence of a *Foxa2, Chrd* and *Noto*-expressing cluster resembling notochord was particularly striking, as prior mouse gastruloid datasets (*11–13*) suggested a complete absence of this cell type. Additionally, notochord differentiation in a human model of trunk development has only recently been achieved under highly specialized culture conditions (*49*). Closer examination of cell line-specific contributions revealed a profound enrichment of *Cnot8* knockout cells in this notochord cluster, alongside a marked depletion from of all other mesodermal cell types and NMPs (Figure 6B, E, F, S7A). In addition, *Cnot8* knockout cells were also overrepresented in undifferentiated epiblast in line with previous reports that *Cnot8* loss interferes with pluripotency exit (*46*). In contrast, endoderm differentiation was preserved or mildly enhanced compared to *Cnot7* knockout and control gastruloids (Figure 6E). *Cnot7* knockout organoids displayed differentiation profiles similar to controls, supporting their utility as a second control condition (Figure 6E, S7A). Differential expression analysis between *Cnot8* knockout cells and control cells echoed the observed cell type changes, with downregulation of paraxial mesoderm markers *Tbx6* and *Meox1* and upregulation of neuronal (*Tubb3*) and notochord (*Foxa2, Sox9, Shh, Nog, Chrd*) markers (Figure 6H, S7D). This coincided with a loss of *Hox* genes (Figure 6H, S7C). Focusing on progenitor populations, we combined epiblast, caudo-lateral epiblast (CLE, enriched in *Cnot7* knockout and nontargeting controls) and node-like clusters (enriched in *Cnot8* knockout), which collectively revealed upregulation of genes involved in neural, notochord and primitive streak differentiation as well as dorso-ventral patterning. In contrast, genes associated with anterior-posterior patterning and mesoderm formation were downregulated (Figure S7E-H).

Further analysis of the putative notochord cluster revealed the presence of notochord-associated signaling factors, including BMP inhibitors *Chrd, Nog*, and *Shh*, as well as the cilia-related genes *Tppp3* and *Fam183b* (Figure 6F-G). This transcriptional profile suggests that these notochord cells represent a functional notochord akin to the embryonic equivalent. Integration of these cells with embryonic data from E7.5-E8.5 embryos (*4*) revealed co-clustering of gastruloid and embryo notochord, with largely overlapping markers (Figure 6K-M), suggesting the presence of *bona fide* notochord cells in *Cnot8* knockout gastruloids.

Given the known function of notochord as a signaling hub in the developing embryo, especially as a source of BMP inhibitors and *Shh*, we compared the signaling landscape of *Cnot8* knockout organoids to *Cnot7* knockout and control. CellChat (*63*) analysis of control cells confirmed the role of NMPs and CLE as a central source of Wnt signaling, particularly of *Wnt3a* (Figure S7I). The depletion of NMPs in *Cnot8* knockouts results in a loss of *Wnt3a* across the organoid. This was also reflected in the downregulation of Wnt target genes, such as *Axin2*, in *Cnot8* knockout organoids and the loss of anterior-posterior patterning markers including *Hox* genes, which are typically expressed at this developmental stage (*11, 12*). Using CellChat, significant Nodal signaling could only be detected in the *Cnot8* knockout organoids, where epiblast, neural crest, endoderm and the notochord itself provided Nodal signals for the notochord (Figure 6N-O). Pseudotime analysis of the combined dataset revealed three independent differentiation pathways starting in the epiblast, with both cell types forming neuronal lineages, but *Cnot8* knockout organoids taking a strikingly altered mesodermal differentiation path, favoring axial over paraxial mesoderm development (Figure 6J).

Together, these data suggest a profound impact of *Cnot8*-mediated deadenylation on lineage specification and body axis formation in gastruloids. Beyond its role in regulating the exit from pluripotency, our scRNA-seq dataset indicates an essential role of *Cnot8* in mesodermal patterning, with *Cnot8* knockout organoids lacking paraxial mesoderm and its derivatives, while gaining node-like and axial mesodermal tissues such as the notochord. This shift completely alters the signaling landscape of *Cnot8* knockout gastruloids, from a Wnt-high to a Wnt-low, *Nodal*- and BMP-inhibitor-rich state (Figure 6P). Our findings establish the transcript poly(A) tail length regulation as a key mediator of embryonic patterning and body planformation.

## Discussion

Here, we leverage the scalability and reproducibility of mouse gastruloids to systematically investigate the role of post-transcriptional regulation in germ layer differentiation. By developing a single-cell CRISPR screening strategy, we identify novel regulators of germ layer commitment. In the endoderm, this includes factors such as *Musashi-2* (*Msi2*), an RNA-binding protein implicated in the regulation of cell proliferation and differentiation in the context of cancer (*64, 65*) and development (*66*). Our data extend *Msi2’s* role to early endodermal lineage specification during mammalian embryogenesis. *Paip2b*, a poly(A)-protein complex binding protein previously implicated in inhibiting cap-dependent translation (*67*), was another candidate showing marked depletion in the NMP compartment. Importantly, depletion of the CCR4-NOT components *Cnot9* and, more prominently, *Cnot8*, but not its paralogue *Cnot7*, results in profound defects in mesoderm differentiation and embryo patterning. These results complement its previously described role in mESC pluripotency exit (*46*) and align with the peri-gastrulation lethality observed in *Cnot8* knockout embryos, a phenotype not seen in *Cnot7* knockout mice, which remain viable (*46*). Given its canonical function as the deadenylase subunit of the CCR4-NOT complex (*51*), the principal deadenylating machinery in the cell, our findings unearth a hitherto unappreciated layer of post-transcriptional control over germ layer specification and body plan formation.

*Cnot8* knockout organoids show dramatic morphological defects, including a lack of axial polarity and sustained *T/Brachyury* expression beyond normal developmental time points. Importantly, we document the emergence of notochord cells starting at 96 h, a cell type absent in mouse gastruloids without modified culture conditions (*49, 60*). These cells express functional signals such as *Shh, Nog*, and *Chrd*, along with cilia-associated genes, suggesting organizer-like activity, thereby reprogramming patterning cues across the entire organoid. Our findings imply a crucial role of *Cnot8* in gastruloid patterning, mediated by its canonical role as deadenylase in poly(A) tail shortening.

Regulation of poly(A) length is crucial for oocyte activation and pre-implantation development, primarily by regulating translational efficiency of mRNAs (*55, 56, 68, 69*). During these early stages, particularly around the transition from the maternal to the zygotic genome, poly(A) tail lengths and transcript stability are decoupled. In contrast, later in development and in mature organisms, poly(A) tail length is no longer thought to influence translation, but instead strongly impacts transcript stability (*68, 69*), with CCR4-NOT-mediated deadenylation promoting transcript destabilization and clearance (*70*). Using Nanopore direct RNA sequencing(*71*), scRNA-seq, ribosome profiling (*72, 73*) and RNA degradation assays, we demonstrate that *Cnot8* loss has negligible effects on translation but rather leads to widespread transcript poly(A) tail elongations and transcript stabilization. This includes transcripts such as *Nodal* (*52*) and *Zic3* (*54, 61*), which are crucial for primitive streak development and its patterning (*53, 54, 61, 62*). Thus, our findings implicate *Cnot8* in regulating the turnover of patterning transcripts beyond the exit from naive pluripotency.

Given *Zic3*’s involvement in node and notochord formation in *Xenopus*(*54*) and mice (*61, 62*), and in left-right symmetry in humans (*74–76*), it is tempting to speculate that *Zic3* represents a strong candidate mediator of the mesodermal patterning defects. In *Xenopus, Zic3* also antagonizes Wnt signaling (*54*), potentially contributing to the diminished Wnt activity observed in *Cnot8*-deficient organoids. The elevated expression of *Zic3* in 120 h progenitor populations supports the hypothesis that *Cnot8* loss may partly phenocopy *Zic3* overexpression, driving aberrant mesodermal patterning. In addition, Nodal signaling is crucial in determining gastruloid patterning (*25, 53*) and a recent study has identified an antagonistic interaction between Wnt signaling and Nodal signaling in establishing the gastruloid body plan (*53*). Ectopic activation of Nodal signaling has been shown to drive early primitive streak characteristics in gastruloids, enabling the differentiation of more anterior, early primitive streak-derived structures, which include the notochord. Although direct *Zic3* overexpression data in mammals are lacking, our results raise the possibility that co-stabilization of *Nodal* and *Zic3*, may be central to the developmental defects seen in *Cnot8* knockouts. This combination of active Nodal and repressed Wnt may drive differentiation of anterior cell lineages typically derived from the early primitive streak (Figure 6P).

Collectively, our findings unveil the regulation of mRNA polyadenylation as a central driver in integrating intrinsic and extrinsic cues to govern mammalian body plan formation and position gastruloid-based single-cell CRISPR screening as a powerful platform for systematic discovery of developmental regulators.

## Acknowledgements

We thank Sendoel lab members for critical input on the manuscript, Catharine Aquino, Anna Bratus-Neuenschwander and the FGCZ for sequencing, Stefanie Jonas for critical discussions and the Cytometry Facility at UZH for assistance with sorting.

## Author contributions

D.T. conducted the experiments and collected the data. N.R and L.L.G assisted with cloning, cell line generation, gastruloid culture, imaging and flow cytometry experiments. D.T., U.G. and A.M. performed bioinformatic data processing and analysis. F.V-F, and P.F.R. assisted with sorting, cloning, virus production and single-cell experiments. C.D. and R.W. assisted with ribosome profiling experiments. K.H. assisted with cloning. S.V. and M.L. assisted with gastruloid culture establishment and experiments. A.M. and M.Z. assisted with Nanopore data analysis. D.T., U.G., A.M. and A.S. performed data analysis and interpretation. D.T. and A.S. conceived the project. A.S. supervised the project.

## Competing interest statement

The authors declare no conflict of interest.

## Materials and Methods

### Mouse embryonic stem cell culture

The dual T/Brachyury-mCherry, SOX1-GFP reporter (SBR) mouse ES cell line was cultured in in tissue culture treated 6-well plates (Thermo Scientific, 140675) (*27*), in DMEM with Glutamax (Gibco, 7001569) supplemented with 10% FCS (2-01F10, BioConcept), Penicillin-Streptomycin (100U/ml, 100µg/ml respectively), non-essential amino acids (0.1 mM, Gibco, 11140050), sodium pyruvate (1mM, Thermo Scientific, 11360070),, beta-mercaptoethanol (0.11 mM, Gibco, 21985023), 3 µM CHIR99021 (SML1046), 1 µM PD0325901 (S1036), 100u/mL LIF (produced in house). Cells were split every 48-72 h by accutase treatment followed by centrifugation at 200 g, 4 min and seeding between 75,000 and 150,000 cells per well.

### Gastruloid culture

Gastruloids were generated as described previously. In short, murine embryonic stem cells (mESCs) are washed twice with PBS, followed by a detachment using Accutase (AT-104) for 4 min at 37°C, 5% CO2. Cells were collected in 4.5 ml of embryonic stem cell medium with 10% fetal calf serum. Cells were centrifuged 200 x g, 4 min, 4°C and then resuspended in 10 ml PBS, followed by a second spin and PBS wash. Finally, cells were resuspended in 1 ml of N2B27 and counted using a hemocytometer. 40 µl of N2B27 with 300 cells per well were seeded in low adhesion 96 well plates (CLS7007-24EA) and incubated for 48 h at 37°C, 5% CO2. After 48 hours, gastruloids were pulsed with 150 µl of N2B27 supplemented with 3 µM CHIR99021 (SML1046-5MG). Medium was refreshed using N2B27 every 24 hours for the extent of the culture (between 120-144 h). For 168 h culture, gastruloids were transferred using a cut tip P1000 pipette into 24-well plates with 500 µl of fresh N2B27 medium and incubated at 37°C, 5% CO2, shaking at 30 rpm on an orbital shaker.

### Cloning of the sgRNA library into the CROP-GFP backbone

669 sgRNA sequences were ordered as IDT oligo pool and cloned in batch using Gibson assembly (HiFi 1 Step Master Mix Kit, SGIDNA Synthetic Genomics GA1100-50) as previously described.(*28, 29*) Following cloning and plasmid purification using the PureLink HiPure Expi Plasmid Megaprep Kit (K210008XP, Thermo), libraries were sequenced to confirm homogeneous sgRNA distribution.

#### Virus production

Vesicular stomatitis virus (VSV-G) pseudotyped lentivirus was produced by calcium phosphate transfection of Lenti-X 293T cells (TaKaRa Clontech, #632180). CROP-Library-GFP construct or CROP-GFP with the individual sgRNAs were transfected together with the plasmids pMD2.G and psPAX2 (Addgene plasmids #12259 and #12260), as previously described.(*28, 29*) Media containing viral supernatant was collected 46 h after initial transfection and passed through a 0.45 µm filter (Sarstedt AG, 83.1826). For the full-scale library, the viral supernatant was further concentrated ∼2,000-fold using a 100 kDa MW cut-off Millipore Centricon 70 Plus (Merck Millipore, UFC710008), the individual guide virus was used without further concentration.

### Murine embryonic stem cell infection – Screen and clonal cell line generation

Before infection for the screen, the virus was titrated by infecting mESCs and measuring GFP signal 3 days post-infection. After identification of a suitable virus dilution to achieve an infectivity of about 10-15%, tet-Cas9 mESCs were infected mixing cells with virus and an infection mix (1/10 dilution of polybrene [10 mg/ml Sigma; 107689-100MG in PBS] in FBS) in a 6 well plate, followed centrifugation at 1100 g for 30 minutes at 37°C. Cells were then cultured for 5 days in standard mESC conditions, followed by induction of Cas9 expression using doxycycline (2µM) for 72h. Cells were then recovered for 24 hours, followed by gastruloid aggregation and single-cell sequencing at 120 h. For clonal cell line generation mESCs were infected with a single-guide carrying virus, aiming for an infectivity of around 10%. Cas9 expression was induced using doxycycline for 72 h, followed by single cell sorting using a BD Aria III cell with a 70 µM nozzle. Clones were genotyped using TIDE assay.(*48*)

### Gastruloid single-cell sequencing

120 h gastruloids were harvested using P1000 cut-tips and subjected to single-cell sequencing. Briefly, gastruloids were collected in 15 ml tubes, washed once with PBS and incubated with accutase for 15 mins at 37°C. Cell suspensions were resuspended using low adhesion tips and centrifuged 200 x g, 4 min, 4°C. For sorting, cells were resuspended in FACS buffer (2% FCS and 0.5 ng/ μl DAPI in PBS), passed through a 35 µm filter cap into 5 ml FACS tubes and sorted for GFP^+^ populations using a BD Aria III cell using a 70 µM nozzle. Sorted cells were then spun down 15 min at 400 x g, 4°C and then resuspended in BD sample buffer. Unsorted gastruloids were harvested and processed the same way, resuspended in sample buffer and then passed through 35 µm cell filters 3 times. For the single-cell CRISPR screen, CROP-GFP+ cells were sorted using a BD Aria III sorter and a 70 µM nozzle. For clonal cell line scRNA-seq, cells were used directly. Single-cell suspensions were prepared for sequencing using the BD Rhapsody Single-Cell Analysis System with the BD Rhapsody Cartridge Kit (633733). Bead-based reverse transcription, amplification and sequencing library production was performed following the manufacturer’s instructions (BD Biosciences, doc ID: 210967 rev. 2.0).

### Targeted sgRNA amplification (Dial-out)

To increase sgRNA annotation in the final dataset, the sgRNA sequence was amplified from BD Rhapsody beads using a nested KAPA PCR, as described previously.(*28, 29*) In the first PCR (PCR1), the Rhapsody beads from individual cartridges were resuspended in 200 ul mastermix (100 µl KAPA HiFi HotStart ReadyMix (Roche 07958935001), 6 µl forward primer (5′-ACACGACGCTCTTCCGATCT-3′, 10µM), 6 µl reverse primer (5′-TCTTGTGGAAAGGACGA-3′, 10µM), 12µl Bead RT/PCR Enhancer reagent from BD Biosciences and 72µl nuclease-free water) and separated into 4 separate 50 µL reactions. Conditions for PCR1: initial denaturation at 95°C / 5 min, 25 cycles of denaturation 95°C / 30 s, annealing 53°C / 30 s, extension 72°C / 20 s and final extension for 10 min at 72°C.

PCR1 products were pooled, followed by bead removal using a magnet and amplicon cleanup by Agencourt AMPure XP beads (Beckman Coulter, A63881) following the manufacturer’s instructions. 3 µl of purified PCR1 product was used as template for PCR 2 and mixed with 47 µl of PCR2 mastermix (25 µL KAPA HiFi HotStart Ready Mix, 2µl forward primer (5′-ACACGACGCTCTTCCGATCT-3′, 10µM), 2 µl reverse primer (CAGACGTGTGCTCTTCCGATCTCTTGTGGAAAGGACGAAACA*C*C* G-3′, 10µM), 18 µl nuclease-free water). PCR2 conditions were: Initial denaturation 95°C / 3 min, followed by 10 cycles of denaturation 95°C / 30 s, annealing 60°C / 3 min, extension 72°C / 60 s, and final extension of 5 min / 72°C. PCR2 product was purified by Agencourt AMPure XP beads and purified PCR2 product was used for indexing PCR. Indexing PCR followed the BD Biosciences ‘mRNA Targeted Library Preparation’ protocol (doc ID: 210968 rev. 3.0).

### Bulk-RNA-seq

RNA was isolated from ribosome profiling lysate buffer lysates using Directzol RNA Kit (Zymo Research, R2052), according to the manufacturer’s instructions. Purified RNA was sent to the Functional Genomics Center Zürich or Novogene for library preparation and sequencing.

#### Ribosome profiling

Ribosome profiling experiments were performed as previously described(*29, 73*) with minor modifications for gastruloids. In short, gastruloids were harvested at indicated time points using cut P1000 tips and collected into 15 ml tubes. Gastruloids were washed 2 times in PBS+CHX, supernatant was removed, and organoids were flash frozen in liquid nitrogen. Frozen organoids were then resuspended in ribosome lysis buffer (1x mammalian polysome buffer, 1 mM DTT, 1% TritonX-100, 0.5% NP40, 25 U/µL Turbo DNase, 100 µg/mL CHX), incubated for 5 min on ice, followed by 5 min spin at 20,000 g at 4°C. Supernatant was collected and flash frozen in liquid nitrogen, before being stored at −80°C before downstream processing.

For ribosome profiling library preparation, samples were thawed and subjected to RNAse digest (1 U/µg of RNase 1 (Epicentre/Lucigen, 10U/µL) for 45 min at room temperature, which was stopped by adding 4 µL of SUPERase Inhibitor (20 U/µL, SUPERase-In RNase Inhibitor Thermo Scientific AM2694). Ribosome-protected footprints were first purified using Amersham MicroSpin S-400 HR columns (Cytiva, 27514001) and then isolated using Direct-zol RNA Kit, according to the manufacturer’s instructions. Total RNA concentration was measured using Qubit RNA broad range kit (Invitrogen, Q10210). Samples were then loaded onto a 15% TBE-UREA gel (Invitrogen, EC68852BOX) and ribosome-protected footprints were excised and extracted using the 20nt, 22nt, 27nt, 30nt and 60nt oligonucleotides as markers (1 µL of 10 µM dilution in 19µL RNase-free H2O). Extraction from gel pieces was performed overnight using RNA extraction buffer (300 mM NaOAc pH=5.5, 1mM EDTA, 0.25% v/v SDS). Extracted footprints were subjected to T4 PNK (NEB, M0201S) end-healing for 1 hour at 37°C. Linker ligation was performed by combining 20 µM of barcoded preadenylated linker with T4 RnI truncated K227Q (NEB, M0351L) for 3 hours at 22°C (50% w/v PEG-8000, 10X T4 RNA ligase, preadenylated linker, T4 RnI truncated K227Q). Samples were then pooled to reduce sample number. This was followed by ribosomal RNA (rRNA) depletion by mixing 10 µL of sample with 10 µL rRNA depletion oligos (2 µM), incubating for 2 minutes at 80°C, followed by a gradual decrease in temperature to 25°C and incubation with MyOne Streptavidin dynabeads (Life Technologies, 65001) to remove rRNA depletion oligos. SuperScript IV Reverse Transcriptase (Invitrogen, 18090050) was used for reverse transcription of rRNA-purified samples. cDNA was then run over a 15% TBE-UREA gel and footprints were excised and extracted using DNA extraction buffer (10mM Tris-HCl pH=8, 300mM NaCl, 1mM EDTA) overnight. Extracted cDNA was circularized using CircLigase II (100 U/µL, Biosearch, CL4115K) for 1 hour at 60°C, followed by PCR amplification using Phusion High-Fidelity DNA polymerase (Thermo Scientific, F530S). After amplification, amplicons between 6 and 12 cycles were visualized using an 8% TBE non-denaturing polyacrylamide gel for the determination of the ideal number of cycles. This was followed by large-scale PCR using barcoded indexing primers and between 7 and 9 cycles. Large scale PCR product was run over an 8% polyacrylamide gel (Invitrogen, EC61252BOX). PCR product was excised from the gel and extracted using DNA extraction buffer by incubating overnight. DNA was then recovered and precipitated using GlycoBlue™ Coprecipitant (1:50,000, 15 mg/mL, Thermo Scientific, AM9516) and resuspended in 10 mM Tris-HCl pH=8. DNA quantification and quality control was performed using Qubit 1X dsDNA High Sensitivity Assay (Invitrogen, Q33230) and Bioanalyzer High Sensitivity DNA kit (Agilent, 5067-4626). Samples were pooled for sequencing.

### Nanopore direct RNA sequencing

Samples for Nanopore direct RNA sequencing were harvested at 72 h of gastruloid development. Gastruloids were harvested using cut P1000 tips, collected in 15 ml tubes and washed 2 times in PBS+CHX. Supernatant was removed, and organoids were flash frozen in liquid nitrogen. Frozen organoids were then resuspended in ribosome lysis buffer (1x mammalian polysome buffer, 1mM DTT, 1% TritonX-100, 0.5%NP40, 25U/µL Turbo DNase, 100µg/mL CHX), incubated for 5 min on ice, followed by 5 min spin at 20,000 g at 4°C. Supernatant was collected and flash-frozen in liquid nitrogen, before being stored at −80° C for downstream processing. RNA was isolated from ribosome profiling lysate buffer lysates using Direct-zol RNA Kit (Zymo Research, R2052), according to the manufacturer’s instructions. Nanopore direct RNA-sequencing library preparation was performed using the Direct RNA Sequencing Kit (SQK-RNA004) according to the manufacturer’s instructions. Samples were loaded onto MinION flow cells (FLO-MIN114).

### RNA degradation experiments

Gastruloids at 96 h of development were treated with 10 µM Actinomycin D (ThermoScientific, 11805017) in N2B27 and gastruloids were harvested after 0, 2 and 4 h of treatment. Gastruloids were harvested at indicated time points using cut P1000 tips and flash frozen. Frozen organoids were then resuspended in ribosome lysis buffer without cyclohexamide (1x mammalian polysome buffer, 1mM DTT, 1% TritonX-100, 0.5%NP40, 25U/µL Turbo DNase), incubated for 5 min on ice, followed by 5 min spin at 20,000 g at 4°C. Supernatant was collected and RNA was purified using the DirectZol RNA kit (Zymo Research, R2052). Purified RNA was submitted for library preparation and sequencing to Novogene.

### Bulk RNA sequencing of sorted cells

T/Brachyury-P2A-mCherry+ cells were sorted using a BD Aria III sorter and a 70 µM nozzle. Sorted cells were spun down and the pellet was suspended in Trizol. RNA was purified using the DirectZol RNA kit, including on-column DNase digest and purified RNA was submitted to Novogene for library preparation and sequencing.

### Immunostaining and imaging

Gastruloids were harvested at indicated time points using cut P1000 tips, washed with PBS and then transferred into 2 ml of 4% paraformaldehyde (PFA) in PBS. Immunostaining was performed as previously described (CITATION: Vianello). Gastruloids were fixed in PFA for 2 hours at 4C, followed by 3 washes in PBS and 2 hours blocking in PBS-FT (PBS with 10% FCS, 0.1% Triton). Primary antibodies were added in PBS-FT and gastruloids were incubated overnight at 4°C, with shaking. This was followed by 3 washes in PBS-FT, followed by incubation with secondary antibody in PBS-FT overnight at 4°C, with shaking. After 3 more washes in PBS-FT, gastruloids were transferred into PBS and imaged using a Zeiss Axio Observer.

### Flow-cytometry

Gastruloids were harvested at the indicated time points using cut P1000 tips and transferred into 15 ml tubes. Media supernatant was removed and gastruloids were washed once with PBS. Gastruloids were resuspended in 500 µl of Accutase and incubated for 10 min at 37°C. Gastruloids were then dissociated using pipetting with low-adherence tips, followed by an additional 5 min of incubation. Cells were then centrifuged 200 x g, 4 min, 4°C, supernatant was removed, and cells were resuspended in FACS buffer (2% FCS and 0.5 ng/ μl DAPI in PBS). DAPI, GFP, and mCherry signal was measured using BD LSRFortessa™ flow cytometer.

### Western blot

TetCas9-SBR mESCs cells were collected from a 6-well plate. After one wash with 1x PBS (Gibco 10010-015), cells were scraped off and lyzed for 10 minutes with RIPA buffer (Sigma R0278) with 1X Protein Cocktail inhibitor (50X Promega, G6521) on ice. Lysate was spun for 10 min at 20,000 g, and supernatant was collected. Protein lysate was mixed with NuPAGE LDS Sample buffer (4X, Invitrogen NP0007) and NuPAGE Sample Reducing Agent (10X, Invitrogen NP0009). Protein samples were denatured at 95°C for 5 minutes and loaded on a 4-12% NuPage 12-well Bis-Tris Gel (Invitrogen, #7001691) using 1X NuPAGE MOPS SDS Running Buffer (20X, Invitrogen, NP0001). Following protein transfer to a nitrocellulose membrane (Cytiva Amersham Protran 0.45 µm NC, 10600002) by tank transfer at 4°C, 30V for 90 minutes, membranes were blocked using 5% skim milk at room temperature for 2 hours. Primary antibody staining was performed overnight at 4°C, followed by 3 washes with TBS-Tween 0.1%. Following the washes, membranes were incubated with secondary antibody for 2 hours at 4°C, followed by 3 additional washes with TBS-Tween 0.1%. Membranes were developed with freshly mixed ECL solutions (Amersham Cytiva. RPN2209). Antibodies used in this study: Cas9 (7A9-3A3) mouse mAb (Cell signaling #1497), Actin (8H10D10) mouse mAb (Thermo Scientific MA5-15452), Cas9 (7A9-3A3) mouse mAb (Cell signaling #1497) and Anti-mouse IgG HRP-linked antibody (Cell signaling #7076).

### Bioinformatic analysis of single-cell sequencing data

Raw sequencing data was processed as previously described.(*28, 29*) In short, sequencing data in BCL format were demultiplexed into FASTQ files using Illumina’s bcl2fastq (v2.20). The analysis was performed with default settings, permitting up to one base mismatch in the sample barcode sequences. The resulting FASTQ files were then processed through the BD Rhapsody™ Analysis Pipeline (v1.9.1), executed on the Seven Bridges cloud genomics platform. For alignment, a custom mouse reference genome was created using Gencode vM25 (GRCm38.p6), which was augmented with the sequences of 720 single guide RNAs (sgRNAs). A custom STAR genome index was generated from this reference. Alignment was performed within the pipeline with the ‘Refined Putative Cell Calling’ option disabled. Following alignment and cell calling, the raw unique molecular identifier (UMI) counts were corrected using the Recursive Substitution Error Correction (RSEC) algorithm provided by BD Genomics to generate the final gene expression matrices for downstream analysis.

### Bioinformatic analysis of dial-out data

PCR dial-out data were processed using an in-house custom Python script as previously described.(*28, 29*) Three 9-nucleotide cell barcodes were extracted from Read 1, allowing for up to one mismatch for each barcode (Hamming distance of 1). The script retrieved the 8-nucleotide UMI sequences from Read 1 as per the BD manual. This was followed by sgRNA annotation in Read 2 for each valid barcode, using an exact match search for the 20-nucleotide sgRNA sequence of the 669 sgRNAs in the screen or the 5 sgRNAs used to demultiplex the individual clonal cell lines in the Cnot7 and *Cnot8* knockout scRNA-seq experiment. Following UMI deduplication, sgRNA UMI counts per cell were computed. In case of multiple sgRNAs detected for an individual cell, specific sgRNA assignment was only performed if the cell’s UMI count exceeded the 99th percentile.

### Single-Cell RNA-seq Data Analysis

Doublets were identified and removed from each sample using the Scrublet Python package(*77*), with expected doublet rates adjusted based on estimates from the BD Rhapsody scanner. For downstream processing, only cells classified as singlets and containing a detected sgRNA were retained. Cells were further filtered based on quality control metrics, requiring UMI counts > 500, UMI counts below the 99th percentile to remove outliers, and mitochondrial gene content < 20%. The filtered dataset was processed using the standard Seurat pipeline. Gene expression was normalized using the NormalizeData function, which scales counts to 10,000 per cell, followed by a natural log transformation. The top 2,000 highly variable genes were identified using FindVariableFeatures, and the data was scaled with ScaleData. Principal Component Analysis (PCA) was performed with RunPCA, and the top 50 principal components were used to construct a nearest-neighbor graph (FindNeighbors). Cell clusters were identified using the FindClusters function with Leiden modularity optimization.(*78*) Data was visualized in two dimensions using Uniform Manifold Approximation and Projection (UMAP).

Batch correction for the screen data was performed using Harmony. Marker genes for each cluster were identified with FindAllMarkers, and clusters were manually annotated based on the expression of canonical cell type markers. Differential expression analysis was conducted using both the MAST package and the Wilcoxon rank-sum test. Kernel density of gene expression was estimated using the Nebulosa package. The dataset was integrated with an external embryo reference using the MouseGastrulationData package(*4*) and the FindTransferAnchors and MapQuery functions in Seurat. Cell signaling pathways were analyzed with CellChat, and pseudotime trajectories were inferred using Monocle3.(*79–81*)

### Bulk RNA Sequencing Data Analysis

Paired-end bulk RNA-seq data was processed using the nf-core/rnaseq pipeline (v3.17.0).(*82*) Briefly, raw read quality was assessed using FastQC, followed by adapter and quality trimming with TrimGalore. Reads were aligned to the Gencode vM25 (GRCm38.p6) reference genome using the STAR aligner. Transcript abundance was quantified using Salmon. An aggregated quality control report for the entire analysis was generated by MultiQC.

### Ribosome Profiling Data Analysis

Ribosome profiling data was processed as previously described.(*29*) Ribosome footprints were extracted from FASTQ files using UMI-tools and a custom regular expression pattern. Footprints with a minimum length of 20 nt were retained. Reads aligning to ribosomal and transfer RNAs (rRNA and tRNA) were removed using Bowtie2 (v2.2.5). Reads remaining after rRNA/tRNA removal were aligned to the Gencode vM25 (GRCm38.p6) reference genome using STAR (v2.7.11a), allowing for one mismatch (-- outFilterMismatchNmax 1) and up to 20 multi-mapped reads (-- outFilterMultimapNmax 20). PCR duplicates were removed based on UMIs using the UMI-tools dedup command and reads mapping to coding sequence (CDS) regions were counted with featureCounts.

Quality control and periodicity analysis were performed with the Ribowaltz package.(*83*) Deduplicated, transcriptome-aligned BAM files were used as input. P-site offsets were calculated using the psite command in auto mode with a flanking region of 6 nucleotides. Standard Ribo-seq quality control plots, including periodicity analysis, were generated using the package’s functions.

### sgRNA enrichment and depletion calculation

The number of sgRNAs in the amplicon sequencing data was counted by the count command of the MAGeCK package.(*40*) To calculate enrichment and depletion of sgRNAs, we used amplicon counts detected in mESCs before doxycycline induction (preDOX) and before gastruloid formation (preGast) compared to the number of cells per sgRNA at 120h. Additionally cell-type-specific depletion and enrichment were computed by comparing cells in one cell type vs all other cells.

### Differential expression

For single-cell differential expression analysis, the DESeq2(*84*) interface with its glmGamPoi R package(*85*) interface was used, which employs a Gamma-Poisson generalized linear model on the data. The p-values were corrected for false discovery rate using the Benjamini-Hochberg procedure. Due to limited cell numbers and statistical power, uncorrected p-values were used for downstream plots in data from the original screen. RNA-seq and Ribo-seq DE and TE analyses were performed by DESeq2 or deltaTE.(*86*)

### Gene ontology analysis

To identify biological processes associated with downregulated genes, Gene Ontology (GO) enrichment analysis was performed. The input gene cutoffs were adjusted to each dataset. The analysis was conducted using the enrichGO function from the clusterProfiler R package (*87*). The function performed an over-representation test on the Biological Process (BP) ontology, using the mouse annotation database (org.Mm.eg.db) with official gene symbols as input. The resulting p-values were adjusted for multiple hypothesis testing. Selected significant (padj < 0.05) GO terms were shown in figures.

### Nanopore data bioinformatic analysis and poly(A) length estimation

Raw nanopore reads in pod5 format were base-called and mapped to the reference mouse genome GRCm38 (primary assembly, obtained from the GENCODE website) using the Dorado software [https://github.com/nanoporetech/dorado] version 0.8.3 with the model rna004_130bps_sup@v5.1.0. Dorado is the software package developed and supported by Oxford Nanopore Technologies for the processing of raw nanopore data obtained on their platform. Dorado uses a deep learning-based algorithm for base-calling (transforming nanopore current data to nucleotide sequence), minimap2 aligner (*88*) for read alignment to the genome, and a separate heuristic algorithm for estimating poly(A) tail length in each read. Importantly, successfully estimated tail lengths are stored as the “pt:i” tags in the output .bam files, while the base-called sequences of reads contain only short fragments of poly(A) tails which are not informative for actual length of the poly(A) tails. Technically, the following command was used:

dorado basecaller rna004_130bps_sup@v5.1.0 $pod5 --estimate-poly-a -- poly-a-config polya_config.toml --mm2-opts “-x splice -Y” --reference GRCm38.primary_assembly.genome.fa

Identification of the poly(A) tails in the reads was configured to tolerate non-adenine gaps up to 3 nt in length: tail_interrupt_length = 3. We further retained only uniquely mapped reads with mapping quality above or equal to one (in minimap2 output, quality=0 indicates that a read is a multimapper that aligns equally well to more than one genomic loci) and present “pt:i” tag. In total, ∼8.1 mln reads were retrieved from four samples (two bioreplicates for each of the two conditions: Cnot8 KO and Control), with ∼1.7 to ∼2.5 mln reads per sample.

To identify the genes from which the reads originated, the following procedure was applied: without considering splicing configuration (“CIGAR” string in the alignment), genomic coordinates of read alignments were intersected with the genomic coordinates of genes from the comprehensive GENCODE annotation of the mouse genome version vM25(*89*) using bedtools intersect version v2.31.1.(*90*) After requiring the minimum overlap to be 0.75 of the read length, ∼5.5 mln reads (∼68%) were assigned to genes. Only ∼230 K reads (∼2.9%) were assigned to more than one gene.

The resulting table of per-read poly(A) tail lengths was further analyzed in R. First, reads were filtered to retain those with a quality score > 10 and a mapping quality > 10. For robust statistical comparison, only genes with at least 50 assigned reads in every sample were included in the differential analysis.

To identify genes with significant changes in poly(A) tail length, a Wilcoxon rank-sum test was performed for each gene, comparing the distributions of individual read poly(A) tail lengths between the Cnot8 KO and control conditions. The effect size was calculated as the log_2_ fold change of the median poly(A) tail length of the KO versus the control. P-values were adjusted for multiple testing using the Benjamini-Hochberg method to calculate the False Discovery Rate (FDR).

### RNA Half-Life Calculation and Statistical Analysis

RNA degradation rates were determined from gene expression data quantified in Counts Per Million (CPM). First, genes with low expression, defined as having a mean of less than 10 CPM across all 0-hour timepoint replicates, were excluded from the analysis. The data normalization was performed in a two-step process. First, to account for initial expression differences, CPM values for each gene at 2 and 4 hours were normalized to the CPM value at the 0-hour timepoint within the same biological replicate. Second, to correct for systemic variations across samples, a scaling factor was calculated for each sample by taking the median of the T0-normalized values from a predefined set of 30 stable housekeeping genes. All T0-normalized expression values for every gene were then divided by their corresponding sample’s scaling factor.

For each gene within each individual replicate, a first-order exponential decay model, N(t)=N0e−kt, was fitted to the fully normalized expression values at 0, 2, and 4 hours using the curve_fit function in the SciPy library. To ensure the reliability of the decay kinetics, fits with a coefficient of determination (R2) below 0.75 were discarded. The RNA half-life (t1/2) was then calculated from the decay rate constant (k) for each valid fit using the formula t1/2=kln(2). To identify significant differences in RNA stability, the calculated half-lives were first log2-transformed. A two-sample Welch’s t-test was performed between the knockout and control groups for each gene with at least two valid half-life values per condition. The resulting p-values were adjusted for multiple comparisons using the Benjamini-Hochberg procedure to control the False Discovery Rate (FDR). Genes with an FDR less than 0.1 were considered to have a statistically significant difference in RNA half-life.

## Supplementary Figures

**Figure S1:**
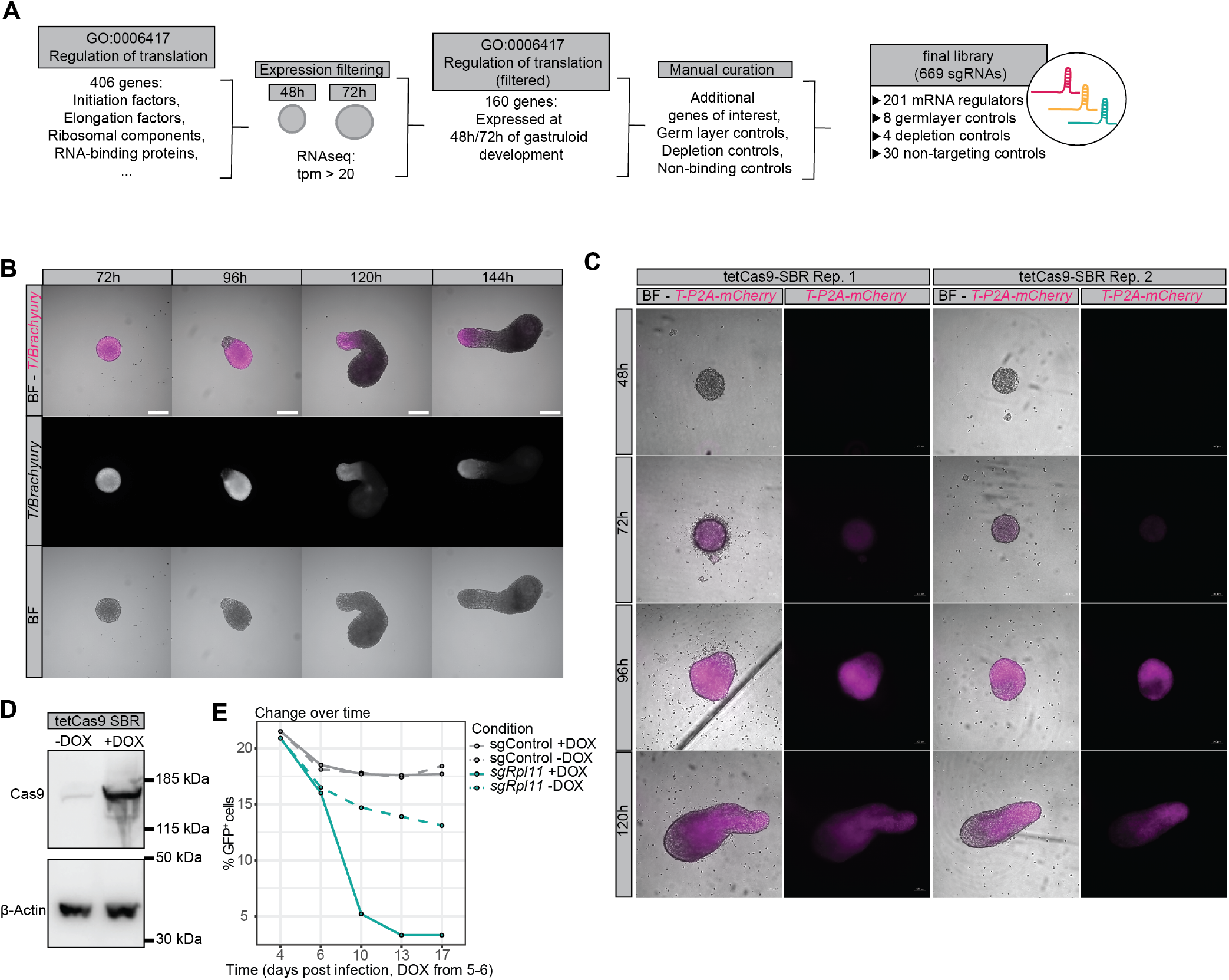
Establishing a tetCas9 inducible murine embryonic stem cell (mESC) line in the SBR background. **(A)** Outline of target gene selection. Genes annotated to the GO term “Regulation of translation (GO:0006417)” were filtered by expression in 48 h and 72 h gastruloids. The resulting list was then manually curated by adding germ layer controls, depletion controls and nontargeting controls were added for a final library of 669 sgRNAs. **(B)** T/Brachyury-mCherry, SOX1-GFP reporter (SBR) mESC-derived gastruloids stained for T/Brachyury at different time points. Scale bar, 200 µM. **(C)** Images of tetCas9 SBR-derived gastruloids during 48, 72, 96 and 120 h of gastruloid development. **(D)** Western blot of CAS9 expression in tetCas9-SBR cells upon 48 h of 2 µM doxycycline treatment. **(E)** Flow cytometry data of tetCas9 SBR cells transduced with sgRNAs targeting *Rpl11* (*sgRPL11*) and nontargeting controls (sgControl), with or without doxycycline induction.

**Figure S2:**
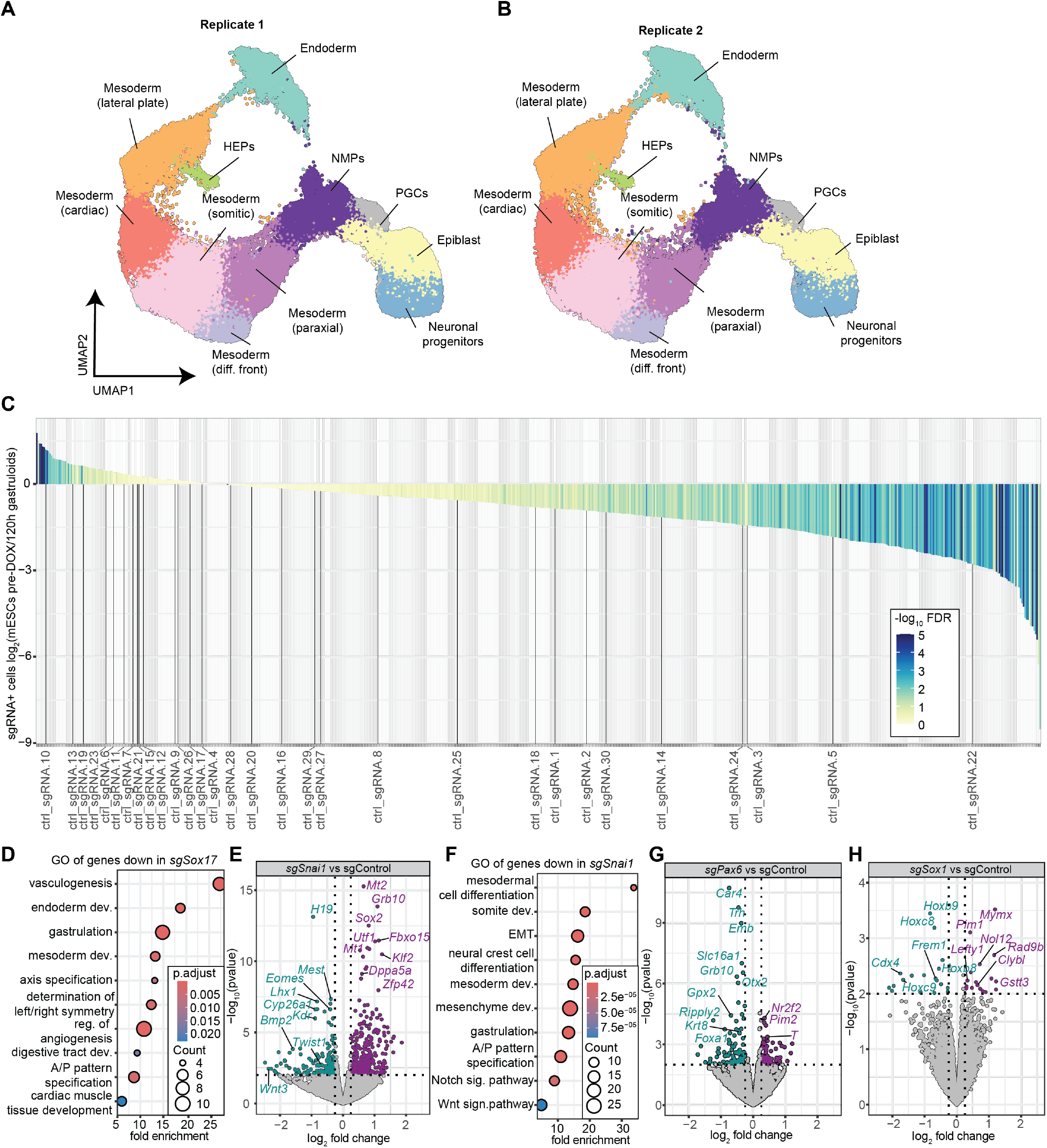
Screen reproducibility and germ layer controls. **(A-B)** UMAP showing cell type distribution of 120 h gastruloids from two independent screen replicates. **(C)** Waterfall plot showing enrichment and depletion of 669 sgRNAs, comparing pre-doxycycline treatment with 120 h gastruloids, with nontargeting control sgRNAs labelled. **(D)** GO term enrichment of genes downregulated (p-value < 0.01) in all *sgSox17* cells. **(E)** Volcano plot displaying the top differentially expressed genes in 120 h gastruloids upon perturbation of *Snai1* (*sgSnai1*). **(F)** GO term enrichment of genes down-regulated (p-value < 0.01) in all *sgSnai1* cells. **(G)** Volcano plot displaying the top differentially expressed genes in 120 h gastruloids upon perturbation of *Pax6* (*sgPax6*). (H) Volcano plot displaying the top differentially expressed genes in 120 h gastruloids upon perturbation of *Sox1* (*sgSox1*).

**Figure S3:**
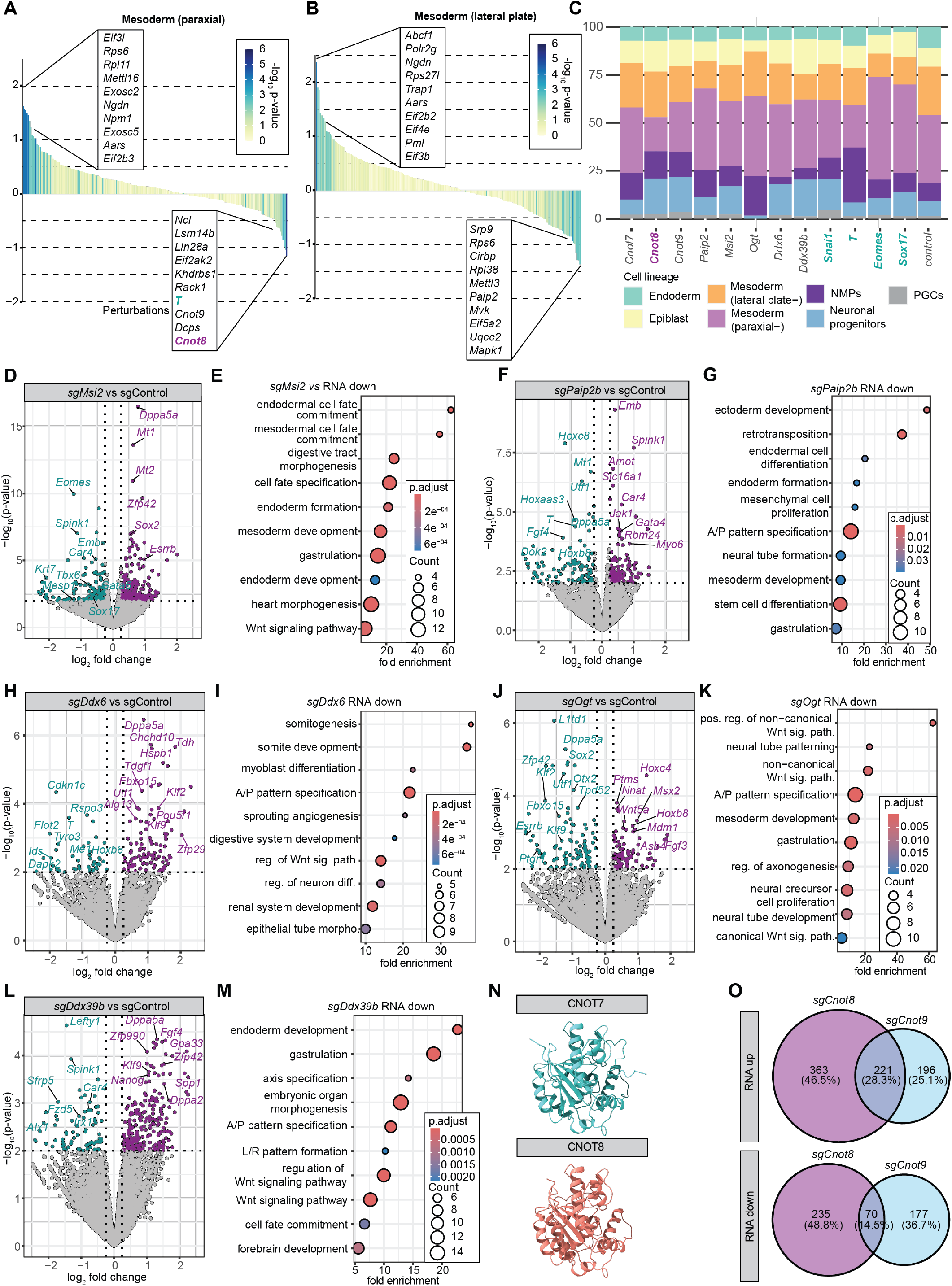
Identification of additional novel regulators of germ layer differentiation in 120 h gastruloids. **(A-B)** Waterfall plots displaying the changes in sgRNA representation in paraxial **(A)** and lateral plate mesoderm **(B). (C)** Higher magnification of Figure 2E, highlighting candidates, germ layer controls and nontargeting controls **(D-M)** Volcano plot and GO term analysis of additional candidates including endodermally depleted *Msi2* (*sgMsi2*) **(D-E)**, neuromesodermally depleted *Paip2b* (*sgPaip2b*) **(F-G)**, globally depleted *Ddx6* (*sgDdx6*) (H-I), neuronally depleted *Ogt* (*sgOgt*) (J-K) and globally depleted *Ddx39b* (*sg Ddx39b*). (L-M). **(N)** Alphafold structure of CNOT7 and CNOT8. **(O)** Venn diagram of genes significantly upregulated (top panel) or downregulated (bottom panel) in *sgCnot8* and *sgCnot9* cells.

**Figure S4:**
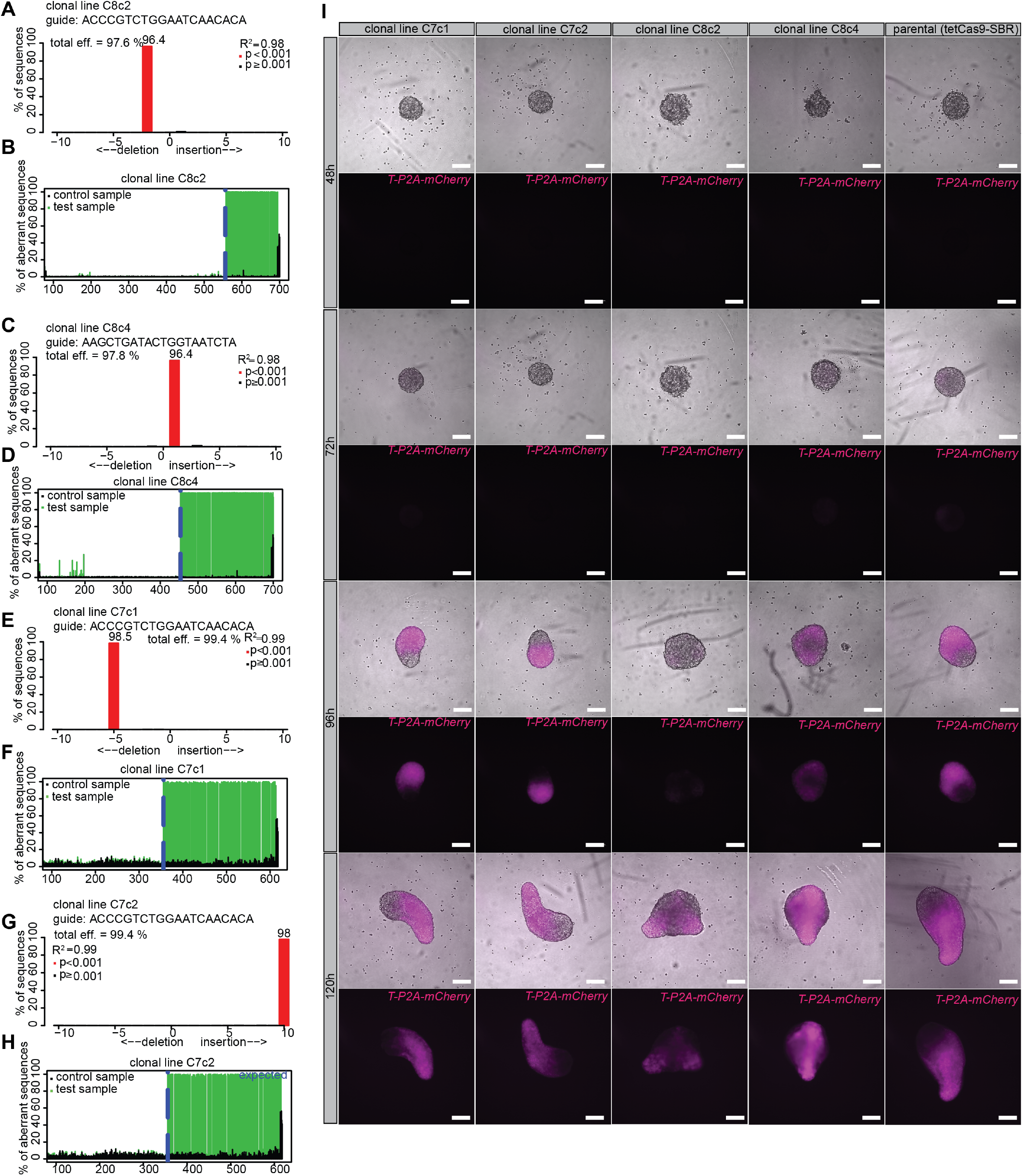
Establishing *Cnot7* and *Cnot8* knockout clonal cell lines. **(A-H)** Tide assay results for 2 *Cnot8* knockout **(A-D)** and 2 *Cnot7* knockout clonal cell lines **(E-H). (I)** Time-lapse images of gastruloids from 2 *Cnot7* clonal cell lines, 2 *Cnot8* clonal cell lines, alongside the parental control lines (tetCas9-SBR).

**Figure S5:**
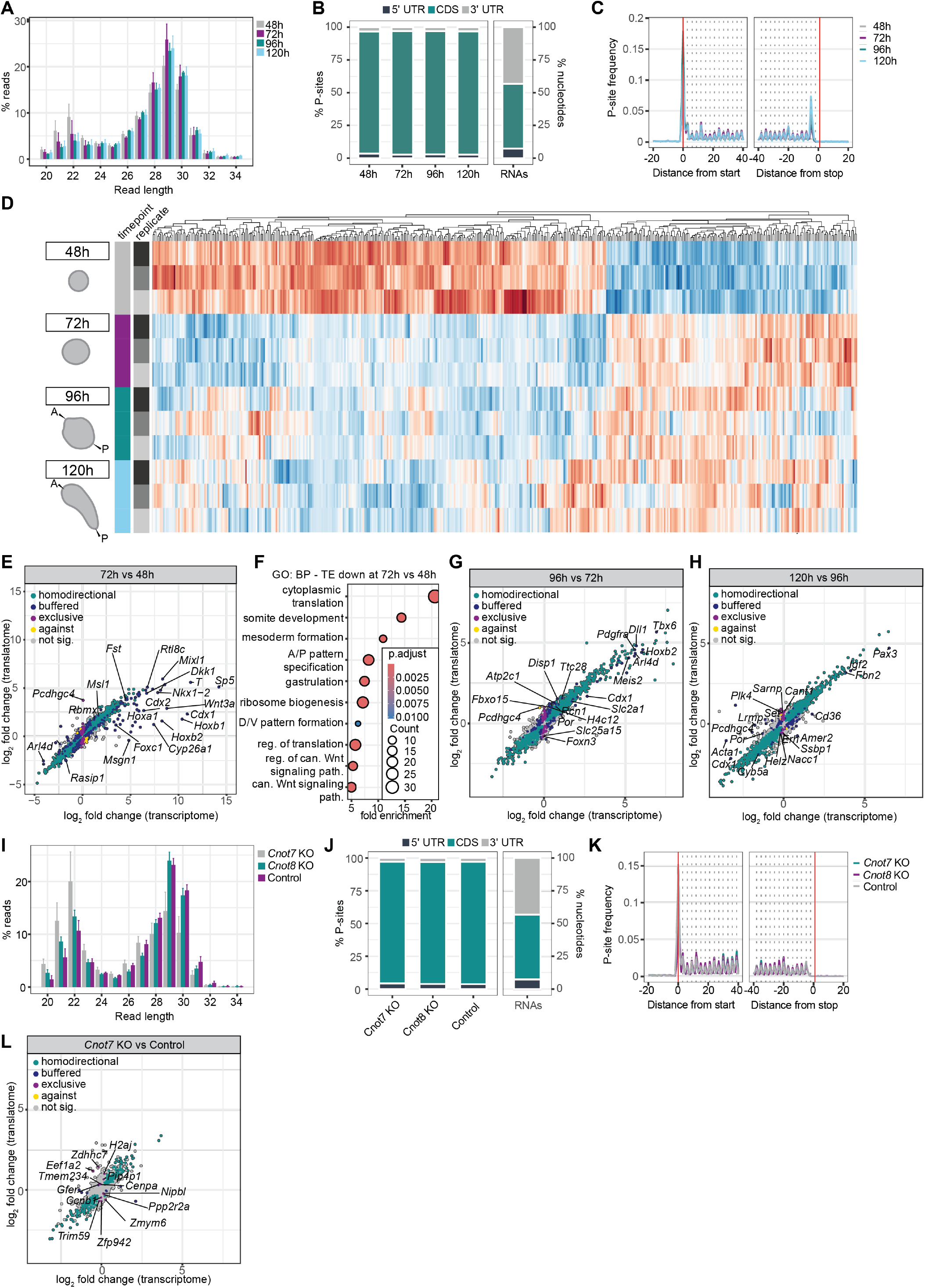
Establishing the translational landscape of gastruloids. **(A)** Fragment length distribution of ribosome profiling samples from 48 h, 72 h, 96 h and 120 h gastruloids. **(B)** P-site distribution of ribosome profiling samples from 48 h, 72 h, 96 h and 120 h gastruloids shows enrichment in the coding sequence (CDS). **(C)** Periodicity plot showing triplet periodicity in ribosome profiling samples from 48 h, 72 h, 96 h and 120 h gastruloids. **(D)** Heatmap showing the translational efficiency (TE) of 483 genes with significantly altered TE between 72 h and 48 h of gastruloid development. **(E-H)** Scatter plot showing the log_2_ fold changes in transcript abundance and ribosome occupancy (translatome) of **(E)** 72 h versus 48 h, **(G)** 96 h versus 72 h **(H)** and 120 h versus 96 h gastruloids. Genes regulated exclusively at the translational level are shown in magenta. (F) GO terms enriched among genes with decreased translational efficiency (TE) at 72 h versus 48 h of gastruloid development. **(I)** Fragment length distribution of ribosome profiling samples from 48 h, 72 h, 96 h and 120 h gastruloids. **(J)** P-site distribution of ribosome profiling samples from *Cnot7* KO, *Cnot8* KO and control gastruloids at 120 h shows enrichment in the coding sequence (CDS). **(K)** Periodicity plot showing triplet periodicity in ribosome profiling samples from *Cnot7* KO, *Cnot8* KO and Control gastruloids at 120h. **(L)** Scatter plot showing the log_2_ fold changes in transcript abundance and ribosome occupancy (translatome) of *Cnot7* knockout vs control gastruloids.

**Figure S6:**
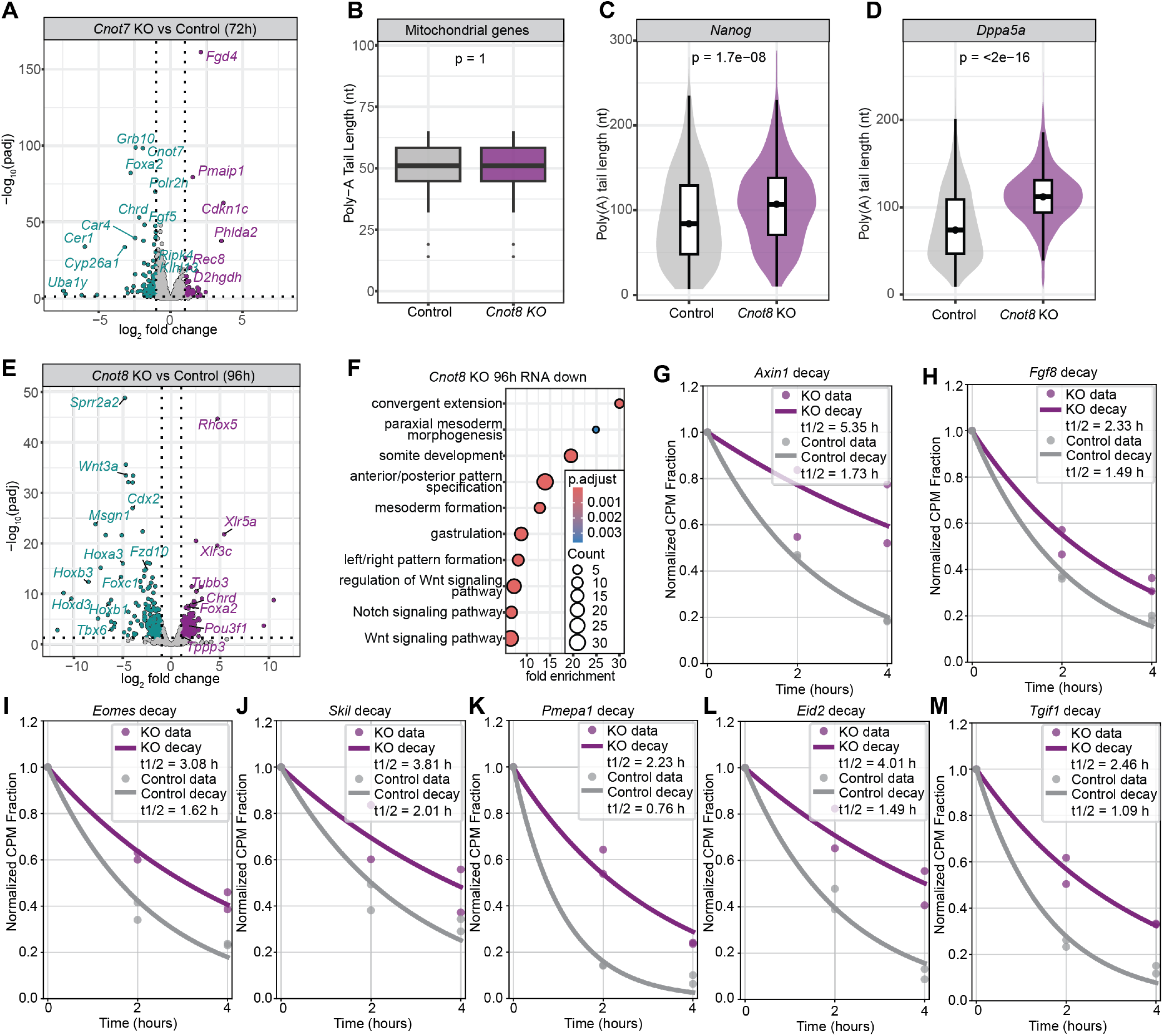
Poly(A) length measurements and RNA degradation. **(A)** Volcano plot displaying the top differentially expressed genes in 72 h *Cnot7* knockout gastruloids versus controls (replicates = 3). **(B)** Box plot of median poly(A) tail lengths of mitochondrial genes. (C-D) Box plot showing poly(A) tail lengths of *Nanog* **(C)** and *Dppa5a* **(D). (E)** Volcano plot displaying the top differentially expressed genes in 96 h *Cnot8* knockout gastruloids versus controls (replicates =2). **(F)** GO term enrichment of genes downregulated (FDR < 0.05, log_2_ fold change > 1) in 96 h *Cnot8* knockout gastruloids. (G-M) Transcript decay plots of stabilized genes in *Cnot8* KO, including *Axin1* **(G)**, *Fgf8* **(H)**, *Eomes* **(I)**, *Skil* **(J)**, *Pmepa1* **(K)**, *Eid2* **(L)** and *Tgif1* **(M)**.

**Figure S6:**
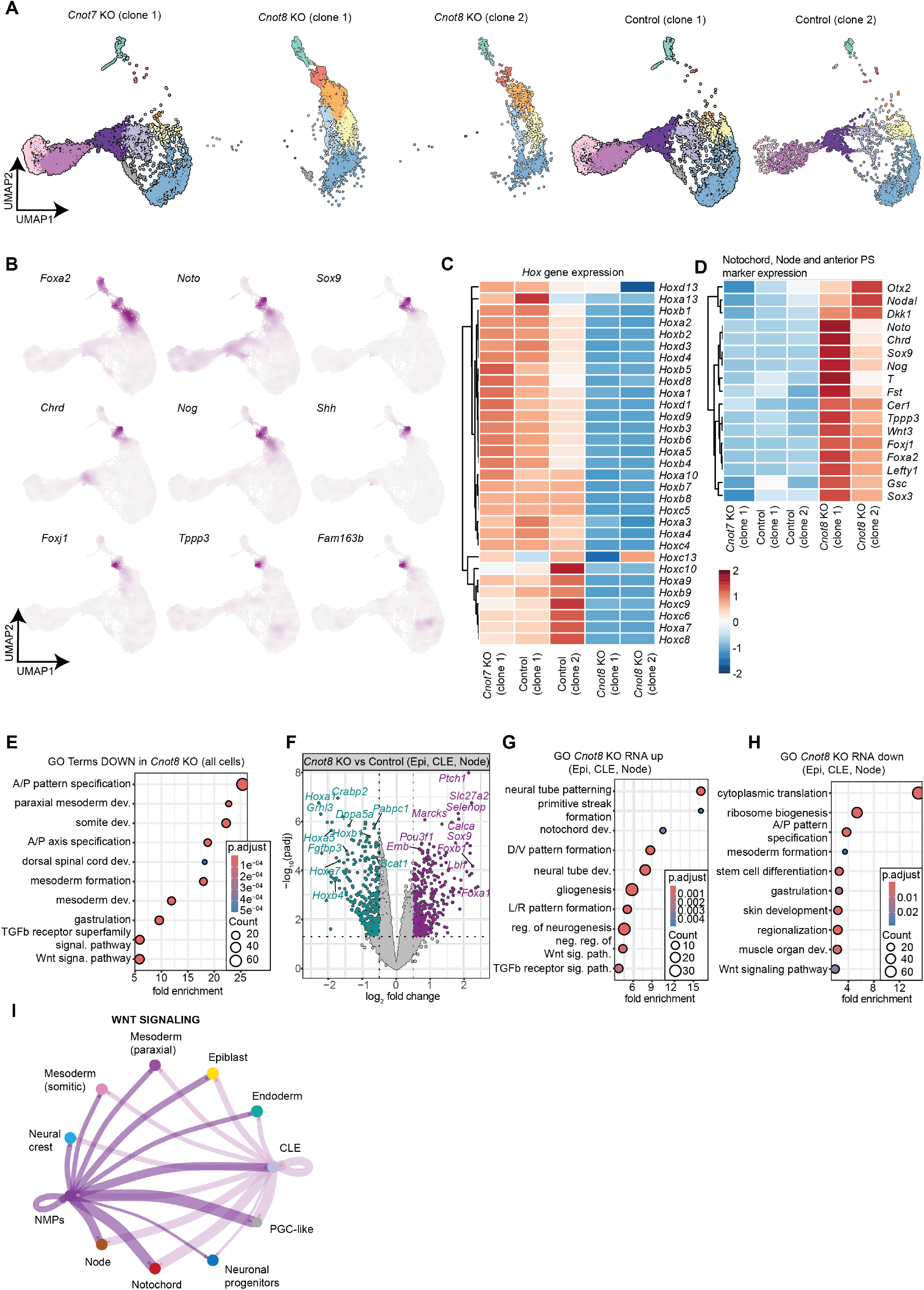
Poly(A) length measurements and RNA degradation. **(A)** UMAP showing cell type distribution of individual clonal cell lines. **(B)** Full-size nebulosa density plots for the notochord markers *Foxa2, Noto, Sox9, Chrd, Nog, Shh, Foxj1, Tppp3* and *Fam183b*. **(C)** Heatmap of *Hox* gene expression in *Cnot7* knockout, *Cnot8* knockout and control gastruloids. (D) Heatmap of markers of the node, the notochord and anterior primitive streak (PS). **(E)** GO term enrichment of genes downregulated (p-adj < 0.05, log_2_ fold change < −1) in all *Cnot8* knockout cells. **(F-H)** Volcano plot **(F)** and GO enrichment analysis **<u>(G-H)</u>** in progenitor clusters (Epiblast, node, caudo-lateral epiblast (CLE)) in *Cnot8* KO cells versus control cells, revealing induction of notochord development and left-right (L/R) pattern formation. **(I)** CellChat analysis of Wnt signaling across the entire dataset (*Cnot7* KO, *Cnot8* KO, control) identifies NMPs and CLE as major signaling sources.

## Notes

### Competing Interest Statement

The authors have declared no competing interest.

## References

1. E. S. Bardot, A.-K. Hadjantonakis, Mouse gastrulation: Coordination of tissue patterning, specification and diversification of cell fate. Mechanisms of Development 163, 103617 (2020).

2. M. Leptin, Gastrulation Movements: the Logic and the Nuts and Bolts. Developmental Cell 8, 305–320 (2005).

3. T. Ichikawa, K. Nakazato, P. J. Keller, H. Kajiura-Kobayashi, E. H. K. Stelzer, A. Mochizuki, S. Nonaka, Live imaging and quantitative analysis of gastrulation in mouse embryos using light-sheet microscopy and 3D tracking tools. Nature Protocols 9, 575–585 (2014).

4. B. Pijuan-Sala, J. A. Griffiths, C. Guibentif, T. W. Hiscock, W. Jawaid, F. J. Calero-Nieto, C. Mulas, X. Ibarra-Soria, R. C. V. Tyser, D. L. L. Ho, W. Reik, S. Srinivas, B. D. Simons, J. Nichols, J. C. Marioni, B. Göttgens, A single-cell molecular map of mouse gastrulation and early organogenesis. Nature 566, 490–495 (2019).

5. S. Grosswendt, H. Kretzmer, Z. D. Smith, A. S. Kumar, S. Hetzel, L. Wittler, S. Klages, B. Timmermann, S. Mukherji, A. Meissner, Epigenetic regulator function through mouse gastrulation. Nature 584, 102–108 (2020).

6. J. B. Black, S. R. McCutcheon, S. Dube, A. Barrera, T. S. Klann, G. Rice, S. S. Adkar, S. H. Soderling, T. E. Reddy, C. A. Gersbach, Master Regulators and Cofactors of Human Neuronal Cell Fate Specification Identified by CRISPR Gene Activation Screens. Cell Rep 33, 108460 (2020).

7. L. G. P. Palma, G. M. Kartha, M. Maqueda, M. Barrero, E. Canton Iglesias, J. Gonzalez Miranda, P. Herrero Molinero, R. Torres-Ruíz, B. Payer, C. Bueno, P. Menendez, L. Espinosa, A. Bigas, An unbiased genomewide screen uncovers 7 genes that drive hematopoietic stem cell fate from mouse embryonic stem cells. Blood, blood.2024027742 (2025).

8. P. Datlinger, A. F. Rendeiro, C. Schmidl, T. Krausgruber, P. Traxler, J. Klughammer, L. C. Schuster, A. Kuchler, D. Alpar, C. Bock, Pooled CRISPR screening with single-cell transcriptome readout. Nature Methods 14, 297–301 (2017).

9. A. Dixit, O. Parnas, B. Li, J. Chen, C. P. Fulco, L. Jerby-Arnon, N. D. Marjanovic, D. Dionne, T. Burks, R. Raychowdhury, B. Adamson, T. M. Norman, E. S. Lander, J. S. Weissman, N. Friedman, A. Regev, Perturb-Seq: Dissecting Molecular Circuits with Scalable Single-Cell RNA Profiling of Pooled Genetic Screens. Cell 167, 1853-1866.e17 (2016).

10. S. C. Van Den Brink, P. Baillie-Johnson, T. Balayo, A.-K. Hadjantonakis, S. Nowotschin, D. A. Turner, A. Martinez Arias, Symmetry breaking, germ layer specification and axial organisation in aggregates of mouse embryonic stem cells. Development 141, 4231–4242 (2014).

11. S. C. Van Den Brink, A. Alemany, V. Van Batenburg, N. Moris, M. Blotenburg, J. Vivié, P. Baillie-Johnson, J. Nichols, K. F. Sonnen, A. Martinez Arias, A. Van Oudenaarden, Single-cell and spatial transcriptomics reveal somitogenesis in gastruloids. Nature 582, 405– 409 (2020).

12. L. Beccari, N. Moris, M. Girgin, D. A. Turner, P. Baillie-Johnson, A.-C. Cossy, M. P. Lutolf, D. Duboule, A. M. Arias, Multi-axial self-organization properties of mouse embryonic stem cells into gastruloids. Nature 562, 272–276 (2018).

13. S. Suppinger, M. Zinner, N. Aizarani, I. Lukonin, R. Ortiz, C. Azzi, M. B. Stadler, S. Vianello, G. Palla, H. Kohler, A. Mayran, M. P. Lutolf, P. Liberali, Multimodal characterization of murine gastruloid development. Cell Stem Cell 30, 867-884.e11 (2023).

14. D. A. Turner, M. Girgin, L. Alonso-Crisostomo, V. Trivedi, P. Baillie-Johnson, C. R. Glodowski, P. C. Hayward, J. Collignon, C. Gustavsen, P. Serup, B. Steventon, M. Lutolf, A. A. Martinez, Anteroposterior polarity and elongation in the absence of extraembryonic tissues and spatially localised signalling in Gastruloids, mammalian embryonic organoids. Development, dev.150391 (2017).

15. P. Baillie-Johnson, S. C. Van den Brink, T. Balayo, D. A. Turner, A. M. Arias, Generation of aggregates of mouse embryonic stem cells that show symmetry breaking, polarization and emergent collective behaviour in vitro. Journal of Visualized Experiments 2015 (2015).

16. S. Vianello, M. P. Lutolf, “In vitro endoderm emergence and self-organisation in the absence of extraembryonic tissues and embryonic architecture” (preprint, BioRxiv, 2020); 10.1101/2020.06.07.138883.

17. J. V. Veenvliet, A. Bolondi, H. Kretzmer, L. Haut, M. Scholze-Wittler, D. Schifferl, F. Koch, L. Guignard, A. S. Kumar, M. Pustet, S. Heimann, R. Buschow, L. Wittler, B. Timmermann, A. Meissner, B. G. Herrmann, Mouse embryonic stem cells self-organize into trunklike structures with neural tube and somites. Science 370, eaba4937 (2020).

18. L. Argiro, C. Chevalier, C. Choquet, N. Nandkishore, A. Ghata, A. Baudot, S. Zaffran, F. Lescroart, Gastruloids are competent to specify both cardiac and skeletal muscle lineages. Nature Communications 15, 10172 (2024).

19. G. Rossi, N. Broguiere, M. Miyamoto, A. Boni, R. Guiet, M. Girgin, R. G. Kelly, C. Kwon, M. P. Lutolf, Capturing Cardiogenesis in Gastruloids. Cell Stem Cell 28, 230-240.e6 (2021).

20. L. Braccioli, T. van den Brand, N. Alonso Saiz, C. Fountas, P. H. N. Celie, J. Kazokaitė-Adomaitienė, E. de Wit, Identifying cross-lineage dependencies of cell-type-specific regulators in mouse gastruloids. Developmental Cell, doi: 10.1016/j.devcel.2025.02.013 (2025).

21. G. Rossi, S. Giger, T. Hübscher, M. P. Lutolf, Gastruloids as in vitro models of embryonic blood development with spatial and temporal resolution. Scientific Reports 12, 13380 (2022).

22. D. Cao, J. Bergmann, L. Zhong, A. Hemalatha, C. Dingare, T. Jensen, L. Cox, V. Greco, B. Steventon, B. Sozen, Selective utilization of glucose metabolism guides mammalian gastrulation. Nature 634, 919–928 (2024).

23. A. Villaronga-Luque, R. G. Savill, N. López-Anguita, A. Bolondi, S. Garai, S. I. Gassaloglu, R. Rouatbi, K. Schmeisser, A. Poddar, L. Bauer, T. Alves, S. Traikov, J. Rodenfels, T. Chavakis, A. Bulut-Karslioglu, J. V. Veenvliet, Integrated molecular-phenotypic profiling reveals metabolic control of morphological variation in a stem-cell-based embryo model. Cell Stem Cell 32, 759-777.e13 (2025).

24. C. Dingare, D. Cao, J. J. Yang, B. Sozen, B. Steventon, Mannose controls mesoderm specification and symmetry breaking in mouse gastruloids. Developmental Cell 59, 1523-1537.e6 (2024).

25. K. S. Stapornwongkul, E. Hahn, P. Poliński, L. Salamó Palau K. Arató, L. Yao, K. Williamson, N. Gritti, K. Anlas, M. Osuna Lopez, K. R. Patil, I. Heemskerk, M. Ebisuya, V. Trivedi, Glycolytic activity instructs germ layer proportions through regulation of Nodal and Wnt signaling. Cell Stem Cell 32, 744-758.e7 (2025).

26. S. Stelloo, M. T. Alejo-Vinogradova, C. A. G. H. van Gelder, D. W. Zijlmans, M. J. van Oostrom, J. M. Valverde, L. A. Lamers, T. Rus, P. Sobrevals Alcaraz, T. Schäfers, C. Furlan, P. W. T. C. Jansen, M. P. A. Baltissen, K. F. Sonnen, B. Burgering, M. A. F. M. Altelaar, H. R. Vos, M. Vermeulen, Deciphering lineage specification during early embryogenesis in mouse gastruloids using multilayered proteomics. Cell Stem Cell 31, 1072-1090.e8 (2024).

27. C. Deluz, E. T. Friman, D. Strebinger, A. Benke, M. Raccaud, A. Callegari, M. Leleu, S. Manley, D. M. Suter, A role for mitotic bookmarking of SOX2 in pluripotency and differentiation. Genes Dev 30, 2538– 2550 (2016).

28. P. F. Renz, U. Ghoshdastider, S. Baghai Sain, F. Valdivia-Francia, A. Khandekar, M. Ormiston, M. Bernasconi, C. Duré, J. A. Kretz, M. Lee, K. Hyams, M. Forny, M. Pohly, X. Ficht, S. J. Ellis, A. E. Moor, A. Sendoel, In vivo single-cell CRISPR uncovers distinct TNF programmes in tumour evolution. Nature 632, 419–428 (2024).

29. F. Valdivia-Francia, U. Ghoshdastider, P. F. Renz, D. Spies, M. Ormiston, K. Hyams, C. Shi, C. Duré, M. Yigit, D. Taborsky, A. Khandekar, R. Weber, H. Yamahachi, S. J. Ellis, A. Sendoel, <em>In vivo</em> single-cell CRISPR screening for microproteins identifies a critical ribosomal component. bioRxiv, 2025.03.17.643322(2025).

30. K. M. Schüle, J. Weckerle, S. Probst, A. E. Wehmeyer, L. Zissel, C. M. Schröder, M. Tekman, G.-J. Kim, I.-M. Schlägl, Sagar, S. J. Arnold, Eomes restricts Brachyury functions at the onset of mouse gastrulation. Developmental Cell 58, 1627-1642.e7 (2023).

31. V. Wilson, R. Beddington, Expression of T Protein in the Primitive Streak Is Necessary and Sufficient for Posterior Mesoderm Movement and Somite Differentiation. Developmental Biology 192, 45–58 (1997).

32. V. Wilson, L. Manson, W. C. Skarnes, R. S. Beddington, The T gene is necessary for normal mesodermal morphogenetic cell movements during gastrulation. Development 121, 877–886 (1995).

33. S. J. Arnold, U. K. Hofmann, E. K. Bikoff, E. J. Robertson, Pivotal roles for eomesodermin during axis formation, epithelium-to-mesenchyme transition and endoderm specification in the mouse. Development 135, 501–511 (2008).

34. C. M. Schröder, L. Zissel, S.-L. Mersiowsky, M. Tekman, S. Probst, K. M. Schüle, S. Preissl, O. Schilling, H. T. M. Timmers, S. J. Arnold, EOMES establishes mesoderm and endoderm differentiation potential through SWI/SNF-mediated global enhancer remodeling. Dev Cell 60, 735-748.e5 (2025).

35. Y. Liu, M. Asakura, H. Inoue, T. Nakamura, M. Sano, Z. Niu, M. Chen, R. J. Schwartz, M. D. Schneider, Sox17 is essential for the specification of cardiac mesoderm in embryonic stem cells. Proceedings of the National Academy of Sciences 104, 3859–3864 (2007).

36. X.-B. Qu, J. Pan, C. Zhang, S.-Y. Huang, Sox17 facilitates the differentiation of mouse embryonic stem cells into primitive and definitive endoderm in vitro. Development, Growth & Differentiation 50, 585–593 (2008).

37. M. Viotti, S. Nowotschin, A.-K. Hadjantonakis, SOX17 links gut endoderm morphogenesis and germ layer segregation. Nature Cell Biology 16, 1146–1156 (2014).

38. A. Cano, M.A. Pérez-Moreno, I. Rodrigo, A. Locascio, M. J. Blanco, M. G. del Barrio, F. Portillo, M. A. Nieto, The transcription factor Snail controls epithelial–mesenchymal transitions by repressing Ecadherin expression. Nature Cell Biology 2, 76–83 (2000).

39. E. A. Carver, R. Jiang, Y. Lan, K. F. Oram, T. Gridley, The mouse snail gene encodes a key regulator of the epithelial-mesenchymal transition. Mol Cell Biol 21, 8184–8188 (2001).

40. W. Li, H. Xu, T. Xiao, L. Cong, M. I. Love, F. Zhang, R. A. Irizarry, J. S. Liu, M. Brown, X. S. Liu, MAGeCK enables robust identification of essential genes from genome-scale CRISPR/Cas9 knockout screens. Genome Biology 15, 554 (2014).

41. P. N. Stoney, A. Yanagiya, S. Nishijima, T. Yamamoto, CNOT7 Outcompetes Its Paralog CNOT8 for Integration into The CCR4-NOT Complex. Journal of Molecular Biology 434, 167523 (2022).

42. R. J. Garriock, R. B. Chalamalasetty, M. W. Kennedy, L. C. Canizales, M. Lewandoski, T. P. Yamaguchi, Lineage tracing of neuromesodermal progenitors reveals novel Wnt-dependent roles in trunk progenitor cell maintenance and differentiation. Development 142, 1628–1638 (2015).

43. B. A. Kinney, A. Al Anber, R. H. Row, Y.-J. Tseng, M. D. Weidmann, H. Knaut, B. L. Martin, Sox2 and Canonical Wnt Signaling Interact to Activate a Developmental Checkpoint Coordinating Morphogenesis with Mesoderm Fate Acquisition. Cell Rep 33, 108311 (2020).

44. F. J. Wymeersch, V. Wilson, A. Tsakiridis, Understanding axial progenitor biology in vivo and in vitro. Development 148, dev180612 (2021).

45. A. Bolondi, B. K. Law, H. Kretzmer, S. I. Gassaloglu, R. Buschow, C. Riemenschneider, D. Yang, M. Walther, J. V. Veenvliet, A. Meissner, Z. D. Smith, M. M. Chan, Reconstructing axial progenitor field dynamics in mouse stem cell-derived embryoids. Developmental Cell 59, 1489-1505.e14 (2024).

46. Y. Quan, M. Wang, C. Xu, X. Wang, Y. Wu, D. Qin, Y. Lin, X. Lu, F. Lu, L. Li, Cnot8 eliminates naïve regulation networks and is essential for naïve-to-formative pluripotency transition. Nucleic Acids Res 50, 4414–4435 (2022).

47. M. F. Pera, J. Rossant, The exploration of pluripotency space: Charting cell state transitions in peri-implantation development. Cell Stem Cell 28, 1896–1906 (2021).

48. E. K. Brinkman, T. Chen, M. Amendola, B. van Steensel, Easy quantitative assessment of genome editing by sequence trace decomposition. Nucleic Acids Research 42, e168–e168 (2014).

49. T. Rito, A. R. G. Libby, M. Demuth, M.-C. Domart, J. Cornwall-Scoones, J. Briscoe, Timely TGFβ signalling inhibition induces notochord. Nature 637, 673–682 (2025).

50. J. Warin, N. Vedrenne, V. Tam, M. Zhu, D. Yin, X. Lin, B. Guidoux-D’halluin, A. Humeau, L. Roseiro, L. Paillat, C. Chédeville, C. Chariau, F. Riemers, M. Templin, J. Guicheux, M. A. Tryfonidou, J. W. K. Ho, L. David, D. Chan, A. Camus, In vitro and in vivo models define a molecular signature reference for human embryonic notochordal cells. iScience 27, 109018 (2024).

51. M. A. Collart, The Ccr4-Not complex is a key regulator of eukaryotic gene expression. Wiley Interdisciplinary Reviews: RNA 7, 438–454 (2016).

52. F. L. Conlon, K. M. Lyons, N. Takaesu, K. S. Barth, A. Kispert, B. Herrmann, E. J. Robertson, A primary requirement for nodal in the formation and maintenance of the primitive streak in the mouse. Development 120, 1919–1928 (1994).

53. A. Dias, P. Pascual-Mas, G. Robertson, G. Torregrosa-Cortés, S. Stelloo, P. Casaní-Galdón, S. Babin, Y. Romaniuk, A. Mayran, A. E. Wehmeyer, J. Garcia-Ojalvo, H. M. McNamara, M. Vermeulen, S. J. Arnold, A. Martinez Arias, Opposing Nodal and Wnt signalling activities govern the emergence of the mammalian body plan. bioRxiv, 2025.01.11.632562 (2025).

54. T. J. Fujimi, M. Hatayama, J. Aruga, Xenopus Zic3 controls notochord and organizer development through suppression of the Wnt/β-catenin signaling pathway. Dev Biol 361, 220–231 (2012).

55. S. W. Eichhorn, A. O. Subtelny, I. Kronja, J. C. Kwasnieski, T. L. Orr-Weaver, D. P. Bartel, mRNA poly(A)-tail changes specified by deadenylation broadly reshape translation in Drosophila oocytes and early embryos. eLife 5 (2016).

56. K. Xiang, J. Ly, D. P. Bartel, Control of poly(A)-tail length and translation in vertebrate oocytes and early embryos. Developmental Cell 59, 1058-1074.e11 (2024).

57. M. Blum, S. J. Gaunt, K. W. Cho, H. Steinbeisser, B. Blumberg, D. Bittner, E. M. De Robertis, Gastrulation in the mouse: the role of the homeobox gene goosecoid. Cell 69, 1097–1106 (1992).

58. R. Hernández-Martínez, S. Nowotschin, L. T. G. Harland, Y.-Y. Kuo, B. Theeuwes, B. Göttgens, E. Lacy, A.-K. Hadjantonakis, K. V. Anderson, Axin1 and Axin2 regulate the WNT-signaling landscape to promote distinct mesoderm programs. [Preprint] (2024). 10.1101/2024.09.11.612342.

59. L. Qiu, Y. Sun, H. Ning, G. Chen, W. Zhao, Y. Gao, The scaffold protein AXIN1: gene ontology, signal network, and physiological function. Cell Communication and Signaling 22, 77 (2024).

60. M. Robles-Garcia, C. Thimonier, K. Angoura, E. Ozga, H. MacPherson, G. Blin, In vitro modelling of anterior primitive streak patterning with human pluripotent stem cells identifies the path to notochord progenitors. Development 151, dev202983 (2024).

61. M. J. Sutherland, S. Wang, M. E. Quinn, A. Haaning, S. M. Ware, Zic3 is required in the migrating primitive streak for node morphogenesis and left-right patterning. Hum Mol Genet 22, 1913–1923 (2013).

62. S. M. Ware, K. G. Harutyunyan, J. W. Belmont, Zic3 is critical for early embryonic patterning during gastrulation. Developmental Dynamics 235, 776–785 (2006).

63. S. Jin, C. F. Guerrero-Juarez, L. Zhang, I. Chang, R. Ramos, C.-H. Kuan, P. Myung, M. V. Plikus, Q. Nie, Inference and analysis of cellcell communication using CellChat. Nature Communications 12, 1088 (2021).

64. D.-W. Yeh, X. Zhao, H. R. Siddique, M. Zheng, H. Y. Choi, T. Machida, P. Narayanan, Y. Kou, V. Punj, S. M. Tahara, D. E. Feldman, L. Chen, K. Machida, MSI2 promotes translation of multiple IRES-containing oncogenes and virus to induce self-renewal of tumor initiating stem-like cells. Cell Death Discovery 9, 141 (2023).

65. S. Wang, N. Li, M. Yousefi, A. Nakauka-Ddamba, F. Li, K. Parada, S. Rao, G. Minuesa, Y. Katz, B. D. Gregory, M. G. Kharas, Z. Yu, C. J. Lengner, Transformation of the intestinal epithelium by the MSI2 RNA-binding protein. Nature Communications 6, 6517 (2015).

66. R. Wang, F. Kato, R. Y. Watson, A. M. Beedle, J. A. Call, Y. Tsunoda, T. Noda, T. Tsuchiya, M. Kashima, A. Hattori, T. Ito, The RNA-binding protein Msi2 regulates autophagy during myogenic differentiation. Life Sci. Alliance 7, e202302016 (2024).

67. J. J. Berlanga, A. Baass, N. Sonenberg, Regulation of poly(A) binding protein function in translation: Characterization of the Paip2 homolog, Paip2B. RNA 12, 1556–1568 (2006).

68. A. O. Subtelny, S. W. Eichhorn, G. R. Chen, H. Sive, D. P. Bartel, Poly(A)-tail profiling reveals an embryonic switch in translational control. Nature 508, 66–71 (2014).

69. Z. Xiong, K. Xu, Z. Lin, F. Kong, Q. Wang, Y. Quan, Q. Sha, F. Li, Z. Zou, L. Liu, S. Ji, Y. Chen, H. Zhang, J. Fang, G. Yu, B. Liu, L. Wang, H. Wang, H. Deng, X. Yang, H. Fan, L. Li, W. Xie, Ultrasensitive Ribo-seq reveals translational landscapes during mammalian oocyte-to-embryo transition and pre-implantation development. Nature Cell Biology 24, 968–980 (2022).

70. R. Parker, H. Song, The enzymes and control of eukaryotic mRNA turnover. Nature Structural and Molecular Biology 11, 121–127 (2004).

71. A. Brouze, P. S. Krawczyk, A. Dziembowski, S. Mroczek, Measuring the tail: Methods for poly(A) tail profiling. Wiley Interdiscip Rev RNA 14, e1737 (2023).

72. N. T. Ingolia, S. Ghaemmaghami, J. R. S. Newman, J. S. Weissman, Genome-Wide Analysis in Vivo of Translation with Nucleotide Resolution Using Ribosome Profiling. Science 324, 218–223 (2009).

73. N. T. Ingolia, L. F. Lareau, J. S. Weissman, Ribosome Profiling of Mouse Embryonic Stem Cells Reveals the Complexity and Dynamics of Mammalian Proteomes. Cell 147, 789–802 (2011).

74. M. Gebbia, G. B. Ferrero, G. Pilia, M. T. Bassi, A. S. Aylsworth, M. Penman-Splitt, L. M. Bird, J. S. Bamforth, J. Burn, D. Schlessinger, D. L. Nelson, B. Casey, X-linked situs abnormalities result from mutations in ZIC3. Nature Genetics 17, 305–308 (1997).

75. A. Mégarbané, N. Salem, E. Stephan, R. Ashoush, D. Lenoir, V. Delague, R. Kassab, J. Loiselet, P. Bouvagnet, X-linked transposition of the great arteries and incomplete penetrance among males with a nonsense mutation in ZIC3. Eur J Hum Genet 8, 704–708 (2000).

76. S. M. Ware, J. Peng, L. Zhu, S. Fernbach, S. Colicos, B. Casey, J. Towbin, J. W. Belmont, Identification and functional analysis of ZIC3 mutations in heterotaxy and related congenital heart defects. Am J Hum Genet 74, 93–105 (2004).

77. S. L. Wolock, R. Lopez, A. M. Klein, Scrublet: Computational Identification of Cell Doublets in Single-Cell Transcriptomic Data. Cell Systems 8, 281-291.e9 (2019).

78. V. A. Traag, L. Waltman, N. J. van Eck, From Louvain to Leiden: guaranteeing well-connected communities. Scientific Reports 9, 5233 (2019).

79. C. Trapnell, D. Cacchiarelli, J. Grimsby, P. Pokharel, S. Li, M. Morse, N. J. Lennon, K. J. Livak, T. S. Mikkelsen, J. L. Rinn, The dynamics and regulators of cell fate decisions are revealed by pseudotemporal ordering of single cells. Nature Biotechnology 32, 381–386 (2014).

80. C. Qiu, J. Cao, B. K. Martin, T. Li, I. C. Welsh, S. Srivatsan, X. Huang, D. Calderon, W. S. Noble, C. M. Disteche, S. A. Murray, M. Spielmann, C. B. Moens, C. Trapnell, J. Shendure, Systematic reconstruction of cellular trajectories across mouse embryogenesis. Nature Genetics 54, 328–341 (2022).

81. X. Qiu, Q. Mao, Y. Tang, L. Wang, R. Chawla, H. A. Pliner, C. Trapnell, Reversed graph embedding resolves complex single-cell trajectories. Nature Methods 14, 979–982 (2017).

82. H. Patel, J. Manning, P. Ewels, M. U. Garcia, A. Peltzer, R. Hammarén, O. Botvinnik, A. Talbot, G. Sturm, nf-core bot, M. Zepper, D. Moreno, P. Vemuri, M. Binzer-Panchal, E. Greenberg, silviamorins, L. Pantano, R. Syme, G. Kelly, L. Sola, F. Hanssen, J. A. F. Yates, G. Lichtenstein, J. Espinosa-Carrasco, rfenouil, L. Zappia, C. Cheshire, E. Miller, marchoeppner, P. Zhou, nf-core/rnaseq: nf-core/rnaseq v3.19.0 Tungsten Turtle, version 3.19.0, Zenodo (2025); 10.5281/zenodo.15631172.

83. F. Lauria, T. Tebaldi, P. Bernabò, E. J. N. Groen, T. H. Gillingwater, G. Viero, riboWaltz: Optimization of ribosome P-site positioning in ribosome profiling data. PLoS Comput Biol 14, e1006169 (2018).

84. M. I. Love, W. Huber, S. Anders, Moderated estimation of fold change and dispersion for RNA-seq data with DESeq2. Genome Biology 15, 550 (2014).

85. C. Ahlmann-Eltze, W. Huber, glmGamPoi: fitting Gamma-Poisson generalized linear models on single cell count data. Bioinformatics 36, 5701–5702 (2021).

86. S. Chothani, E. Adami, J. F. Ouyang, S. Viswanathan, N. Hubner, S. A. Cook, S. Schafer, O. J. L. Rackham, deltaTE: Detection of Translationally Regulated Genes by Integrative Analysis of Ribo-seq and RNA-seq Data. Curr Protoc Mol Biol 129, e108 (2019).

87. S. Xu, E. Hu, Y. Cai, Z. Xie, X. Luo, L. Zhan, W. Tang, Q. Wang, B. Liu, R. Wang, W. Xie, T. Wu, L. Xie, G. Yu, Using clusterProfiler to characterize multiomics data. Nature Protocols 19, 3292–3320 (2024).

88. H. Li, Minimap2: pairwise alignment for nucleotide sequences. Bioinformatics 34, 3094–3100 (2018).

89. J. M. Mudge, S. Carbonell-Sala, M. Diekhans, J. G. Martinez, T. Hunt, I. Jungreis, J. E. Loveland, C. Arnan, I. Barnes, R. Bennett, A. Berry, A. Bignell, D. Cerdán-Vélez, K. Cochran, L. T. Cortés, C. Davidson, S. Donaldson, C. Dursun, R. Fatima, M. Hardy, P. Hebbar, Z. Hollis, B. T. James, Y. Jiang, R. Johnson, G. Kaur, M. Kay, R. J. Mangan, M. Maquedano, L. M. Gómez, N. Mathlouthi, R. Merritt, P. Ni, E. Palumbo, T. Perteghella, F. Pozo, S. Raj, C. Sisu, E. Steed, D. Sumathipala, M.-M. Suner, B. Uszczynska-Ratajczak, E. Wass, Y. T. Yang, D. Zhang, R. D. Finn, M. Gerstein, R. Guigó, T. J. P. Hubbard, M. Kellis, A. Kundaje, B. Paten, M. L. Tress, E. Birney, F. J. Martin, A. Frankish, GENCODE 2025: reference gene annotation for human and mouse. Nucleic Acids Res 53, D966–D975 (2025).

90. A. R. Quinlan, I. M. Hall, BEDTools: a flexible suite of utilities for comparing genomic features. Bioinformatics 26, 841–842 (2010).

